# The RecA-directed recombination pathway of natural transformation initiates at chromosomal replication forks in *Streptococcus pneumoniae*

**DOI:** 10.1101/2022.08.04.502747

**Authors:** Calum Johnston, Rachel Hope, Anne-Lise Soulet, Marie Dewailly, David De Lemos, Patrice Polard

## Abstract

Homologous recombination (HR) is a crucial mechanism of DNA strand exchange that promotes genetic repair and diversity in all kingdoms of life. Bacterial HR is driven by the universal recombinase RecA, assisted by dedicated mediators that promote its polymerization on single-stranded DNA (ssDNA). In bacteria, natural transformation is a prominent HR-driven mechanism of horizontal gene transfer specifically dependent on the conserved DprA recombination mediator. Transformation involves internalisation of exogenous DNA as ssDNA, followed by its integration into the chromosome by RecA-directed HR. How DprA-mediated RecA filamentation on transforming ssDNA is spatiotemporally coordinated with other cellular processes remains unknown. Here, we tracked the localisation of functional fluorescent fusions to DprA and RecA in *Streptococcus pneumoniae* and revealed that both accumulate in an interdependent manner with internalised ssDNA at replication forks. In addition, dynamic RecA filaments were observed emanating from replication forks, even with heterologous transforming DNA, which probably represent chromosomal homology search. In conclusion, this unveiled interaction between HR transformation and replication machineries highlights an unprecedented role for replisomes in anchoring transforming ssDNA to the chromosome, which would define a pivotal early HR step for its chromosomal integration.

## Introduction

Homologous recombination (HR) is a universal DNA strand exchange mechanism, which is vital to genome biology *via* its implication in specific pathways of DNA repair and genetic diversification^1–4^. The widely conserved recombinases of the RecA/Rad51 family are core HR effectors that form dynamic nucleofilaments to promote exchange between complementary DNA sequences^5^. These reactions are controlled and assisted by specific effectors, which define different HR pathways across all kingdoms of life. Any dysfunction in these HR assistants can alter cell development, threaten the integrity or adaptive capacity of the genome and endanger cell survival^1, 6, 7^.

Natural transformation is a programmed HR-directed horizontal gene transfer mechanism that is widespread in bacteria and promotes the shuffling of chromosomally-encoded genetic information^8^. As such, transformation facilitates adaptive responses to stresses, including the acquisition of new genetic traits such as antibiotic resistance and vaccine escape ^8–10^, as well as limiting the genetic drift of species by curing genomes of mobile genetic elements ^11–14^.

Transformation is a multistep DNA processing mechanism directed by proteins encoded by the recipient cell (Figure 1A). Most of these are expressed during a distinct physiological state, defined as competence, which is triggered and regulated in different ways depending on the species^8^. Transformation proteins first direct the uptake of exogenous double-stranded DNA (dsDNA) through the cell envelope to the periplasmic space ^15–18;^ next, they couple transport of a linear single-stranded DNA (ssDNA) strand across the cell membrane with degradation of its complementary strand ^16, 19–22;^ internalised ssDNA is then integrated into the genome by RecA-directed HR at homology sites. A key conserved early effector of the HR pathway of transformation is the ssDNA-binding protein DprA, which specifically interacts with RecA to mediate its loading onto ssDNA ^23–25^. Next, as in all HR pathways, RecA polymerises on the ssDNA to form a nucleofilament, referred to as the presynaptic filament, and promotes homology search in chromosomal DNA and pairing with a complementary DNA strand to generate a 3-stranded DNA molecule, defined as the HR heteroduplex, synapse or D-loop. Next, an helicase involved in extending ssDNA incorporation in the genome from the D-loop, which differs from one species to another. In firmicutes, this HR motor is RadA, a protein which also acts with RecA in pathways of DNA repair ^26, 27^. In contrast, a transformation-dedicated helicase ComM is conserved in all other bacterial species^28^. The final reactions of transformation, including covalent linkage and integration of the paired ssDNA molecule with the recipient chromosomal dsDNA, remain uncharacterised. Ultimately, a replication cycle generates a wildtype and a transformed chromosome, each segregated into a daughter cell.

**Figure 1:**
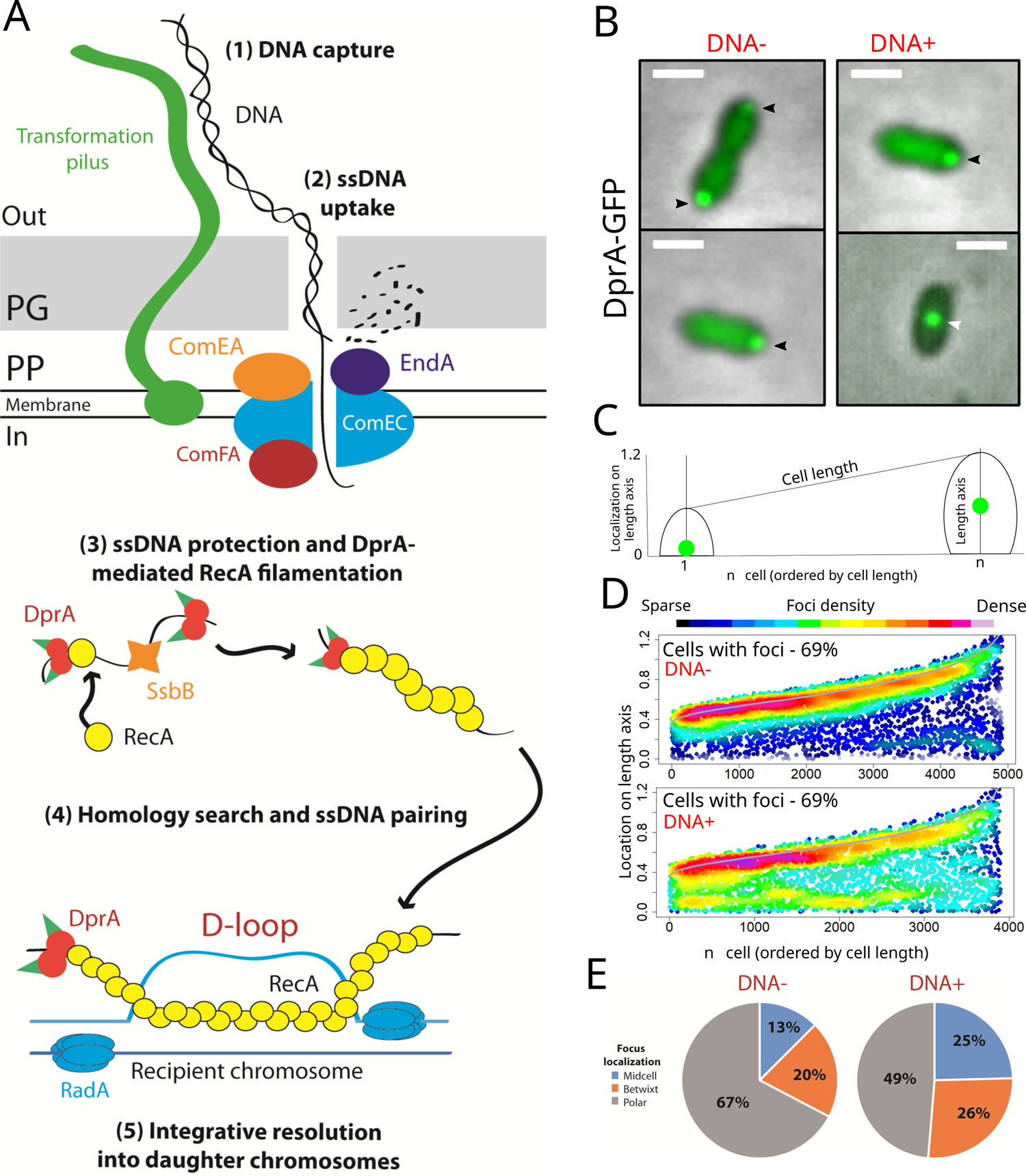
DprA and homologous recombination during transformation. (A) Schematic representation of the steps involved in pneumococcal transformation (1) DNA capture. DNA is captured by a long transformation pilus, formed from the ComG proteins, and transferred to the DNA receptor ComEA. (2) ssDNA uptake. ComEA transfers the DNA to EndA, which degrades one strand of DNA, with the remaining single strand pulled through the ComEC transformation pore by the ComFA ATPase. (3) ssDNA protection and DprA-mediated RecA filamentation. Once internalised, ssDNA interacts with SsbB and the RMP DprA, which loads RecA onto the DNA. Polymerization of RecA along the ssDNA generates the early HR intermediate known as the presynaptic filament. (4) Homology search and ssDNA pairing. The presynaptic filament interacts with the chromosome in an unknown manner, and RecA promotes homology search. Once homology is found, the homologous strand of the recipient chromosome is displaced, and RecA facilitates pairing between the transforming ssDNA and the complementary strand, forming the so-called displacement loop (D-loop). D-loop extension is facilitated by the helicase RadA which unwinds the recipient chromosome on either side of the D-loop. (5) Integrative resolution into daughter chromosomes. The D-loop structure is resolved by the passage of the replication machinery, generating one transformed and one untransformed daughter chromosome. (B) Sample fluorescence microscopy images of R3728 strain (*comC0, dprA-gfp*) producing DprA-GFP 15 minutes after competence induction and 5 minutes after DNA addition (250 ng µL^-1^). Scale bars, 1 µm. Black arrows, polar DprA-GFP foci; white arrows, midcell DprA-GFP foci. (C) Schematic representation of focus density maps with half cells represented as vertical lines in ascending size order and localisation of foci represented along the length axis of each half cell. (D) Addition of transforming DNA shifts the localisation profile of DprA-GFP foci towards midcell. Data represented as focus density maps plotted on the longitudinal axis of half cells ordered by cell length. Each spot represents the localisation of an individual focus, and spot colour represents focus density at a specific location on the half cell. Cells with >0 foci shown for each time point. In cells possessing >1 foci, foci were represented adjacently on cells of the same length. DNA-, 5,739 cells and 4,920 foci analysed; DNA+, 3,406 cells and 3,899 foci analysed. (E) localisation of DprA-GFP foci split into three categories on a half-cell of arbitrary length 1 where midcell is 0 and the pole is 1. Midcell, 0-0.3; betwixt, 0.3-0.7; polar, 0.7-1.

Our current understanding of the transformation mechanism results from studies conducted in a dozen distinct species, including the historical models *Bacillus subtilis* and *Streptococcus pneumoniae* (the pneumococcus), as well as many other human pathogens such as *Haemophilus influenzae, V. cholerae* and *Helicobacter pylori* (for reviews, see ^8, 29^). These studies highlighted important general features of transformation, including the remarkable speed at which transforming DNA (tDNA) is captured, internalised and integrated into the chromosome. This was shown to occur in a minute time frame in *S. pneumoniae* and *V. cholerae*^30, 31^. How the HR system of transformation achieves such efficiency is unexplained. Pioneering studies in *B. subtilis* reported the gradual and stable accumulation of GFP fusions to transformation proteins involved in tDNA uptake and ssDNA transport, as well as RecA, at one pole of competent cells independent of tDNA addition ^32, 33^. In the presence of tDNA, polar RecA evolved into filaments proposed to represent presynaptic filaments formed during the polar entry of ssDNA, which next scan chromosomal DNA for homology ^33, 34^.

Here, we investigated DprA and RecA localisation dynamics during transformation in *S. pneumoniae*. In stark contrast to *B. subtilis*, in which competence occurs in non-replicating cells and lasts for several hours, pneumococcal competence occurs during the exponential phase of growth for a short period of about 30 minutes^35^. In addition, tDNA is captured and enters competent *S. pneumoniae* cells not at the pole but at midcell^36, 37^. Using functional fluorescent fusions of DprA and RecA, we tracked the early HR intermediates of transformation in actively transforming pneumococcal cells. Both proteins formed distinct foci at midcell in transforming cells, dependent on their physical interaction, showing that these nucleoprotein assemblies represent early HR intermediates of transformation. Furthermore, DprA and RecA foci were proven to localise to chromosomal replication forks. Importantly, RecA was observed to form short, dynamic filaments emanating from this replisomal accumulation point, possibly revealing homology search on the chromosome. These results represent an unprecedented link between the HR machinery of natural transformation and the chromosomal replication apparatus, shedding light on the mechanism of targeted homology search on the chromosome during pneumococcal transformation.

## Results

### DprA accumulates at midcell during transformation in S. pneumoniae

To observe the early DprA-mediated HR steps of natural transformation in individual living competent pneumococcal cells, we tracked the localisation of a fluorescent DprA-GFP fusion proven to be fully functional in transformation assays^38^. Purified DprA-GFP was as efficient as DprA in assisting RecA-directed HR in an *in vitro* D-loop assay (Extended Figure 1), validating use of this fusion for analysing DprA localisation dynamics during transformation. We previously showed that DprA-GFP accumulated at a single cell pole in competent cells^38^. This localisation was related to an additional role for DprA in shut-off of pneumococcal competence. This negative feedback loop is independent of the ability of competent cells to uptake tDNA but dependent on a high cellular concentration of DprA^38, 39^. Here, we analysed DprA-GFP localisation upon addition of tDNA to competent, transformable pneumococcal cells. Competence was induced by incubating cells with saturating levels (100 ng mL^-1^) of synthetic competence-stimulating peptide (CSP) for 10 minutes^40^, ensuring all cells in the population were competent. Addition of saturating levels of tDNA (250 ng µL^-1^) then ensured all cells were engaging in transformation. Cells were visualised 5 minutes after tDNA addition and compared with cells without tDNA. DprA-GFP formed foci in competent cells, irrespective of the addition of tDNA (Figure 1B). The frequency and cellular localisation of DprA-GFP foci, presented as focus density maps ordered by cell length (Figure 1C), showed that addition of tDNA did not modify the frequency of foci in competent cells but slightly altered their localisation (Figure 1D), with a significant increase of midcell foci from 13 % to 25 % (Figure 1E). These results suggested that DprA-GFP may interact with internalised ssDNA to generate midcell foci.

We showed previously using a strain expressing *dprA* under the control of an IPTG-inducible P*_lac_* promoter (*CEP_lac_-dprA*) and lacking native *dprA* that reducing cellular levels of DprA in competent cells prevented competence shut-off from occurring whilst maintaining optimal transformation efficiency^39^. Using a *CEP_lac_-dprA-gfp* fusion, we also showed that in similar conditions (6 µM IPTG), no polar foci of DprA-GFP were observed^38^. We took advantage of this strain and these conditions to visualise DprA-GFP during transformation as described above, with 48% of cells possessing DprA-GFP foci, mostly at midcell (Figures 2AB). By contrast, only 7% of cells possessed foci in the absence of tDNA, in agreement with previous results^38^. Most cells present DprA-GFP foci at midcell, while late dividing cells present foci at the ¼ and ¾ positions, future sites of midcell in daughter cells (Figure 2B). To explore where these tDNA-dependent DprA-GFP foci localise along the lateral axis of the cells, data was represented as heatmaps split into six cell categories. tDNA-dependent DprA-GFP foci were present near the centre of the longitudinal axis in all cell types (Figure 2C). In non-constricted cells they were either side of the central axis, while in constricted cells they appeared more central. Thus, DprA was found to accumulate at midcell in a tDNA dependent manner. We next analysed this localisation at different time-points after competence induction and tDNA addition. The results showed that the highest number of cells possessing tDNA-dependent DprA-GFP foci were observed 20 minutes after CSP addition, and that the majority of cells with foci possess a single focus (Extended Figure 2A). In addition, the localisation profile of these foci remains similar over time (Extended Figure 2B). Transforming cells with the same concentration of heterologous chromosomal DNA from *Escherichia coli* resulted in the formation of DprA-GFP foci at a similar frequency and localisation (Figure 2BC), showing that homology between tDNA and chromosomal DNA is not required for focus formation. We next examined how exogenous tDNA concentration impacted DprA-GFP focus formation. Results showed that a 1,000-fold reduction in tDNA concentration, starting from saturating conditions (250 ng µL^-1^), reduced the frequency of cells exhibiting midcell DprA- GFP foci from 47 % to 17 % (Extended Figure 2C). This suggests that the more ssDNA enters each cell, the more DprA-GFP molecules accumulate at midcell to generate detectable fluorescent foci. In conclusion, pneumococcal DprA accumulates at two distinct locations in competent cells, correlating with its two roles in competence and transformation. First, as reported previously^38^, the majority of DprA accumulates at one cell pole to mediate competence shut-off. Second, as observed here, a minority of DprA accumulates at midcell in a tDNA-dependent manner. This clustering of DprA at midcell appears therefore to be related to its role in transformation.

**Figure 2:**
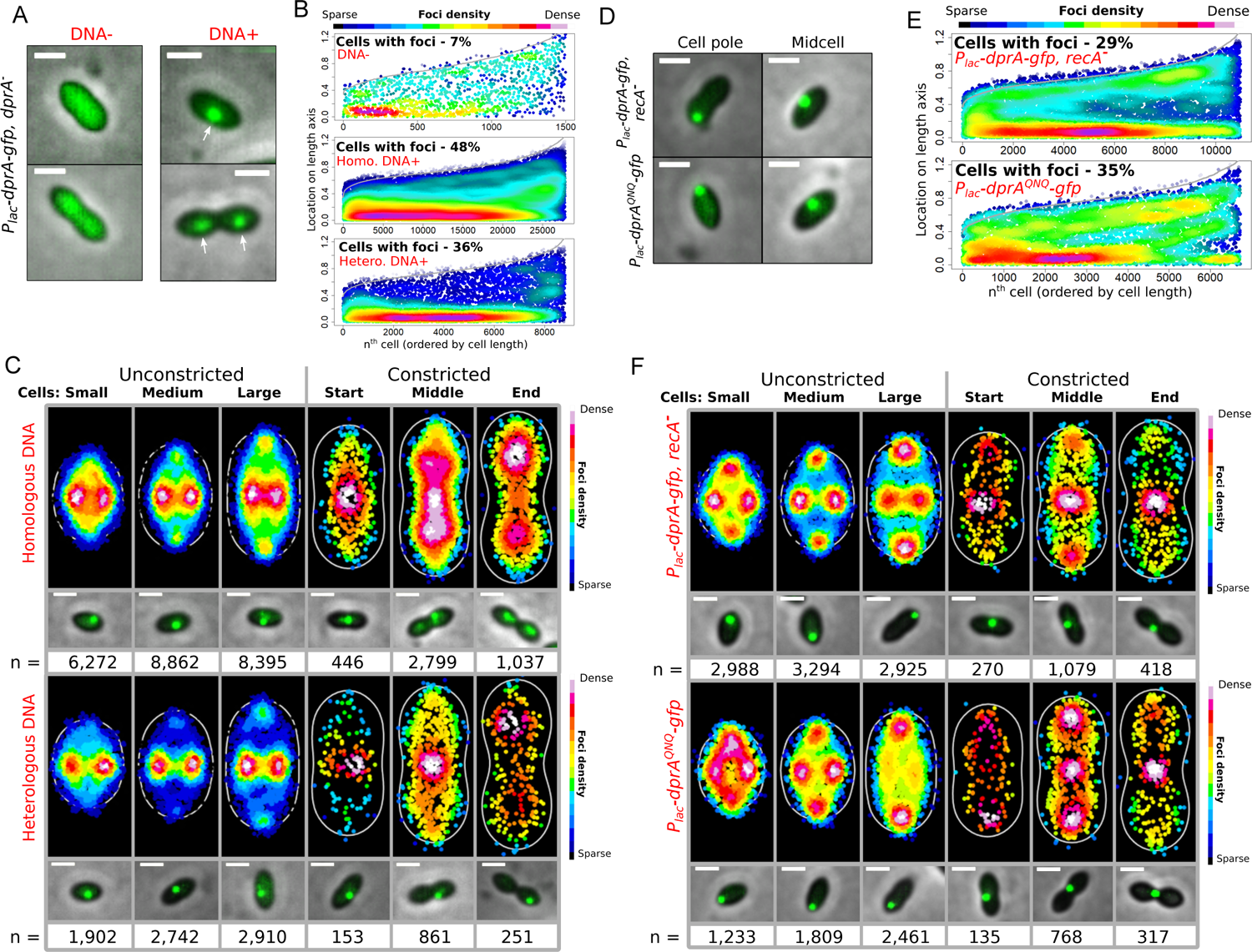
When produced at low levels, DprA-GFP accumulates at midcell upon addition of transforming DNA, dependent on interaction with RecA. (A) Sample fluorescence microscopy images of R4262 strain (*comC0, CEP_lac_-dprA-gfp*, *dprA::spc*) grown in 6 µM IPTG to produce ∼300 DprA-GFP dimers, 15 minutes after competence induction and 5 minutes after DNA addition (250 ng µL^-1^). Scale bars, 1 µm. White arrows, midcell DprA-GFP foci. (B) Low level DprA-GFP accumulates at midcell upon addition of transforming DNA. Representations as focus density maps as described in Figure 1C. DNA-, 21,055 cells and 1,512 foci analysed; Homologous DNA, 54,058 cells and 27,811 foci analysed; Heterologous DNA, 23,962 cells and 8,819 foci analysed. (C) Foci localisation on density heat maps with cells split into six cell cycle categories by sise and constriction status (see Materials and Methods). White lines represent the average cell contour of the sample set and each spot represents an individual focus, with colour representing density. Microscopy images represent sample images of each cell category showing preferential focus localisation. Scale bars, 1 µm. Homologous transforming DNA; Small cells, 17,888 cells and 6,272 foci analysed; medium cells, 16,744 cells and 8,862 foci analysed; large cells, 13,659 cells and 8,395 foci analysed; cons. start cells, 880 cells and 446 foci analysed; cons. middle cells, 3824 cells and 2,799 foci analysed; cons. end cells, 1063 cells and 1,037 foci analysed. Heterologous transforming DNA; Small cells, 7,557 cells and 1,902 foci analysed; medium cells, 7,562 cells and 2,742 foci analysed; large cells, 6,318 cells and 2,910 foci analysed; cons. start cells, 377 cells and 153 foci analysed; cons. middle cells, 1,708 cells and 861 foci analysed; cons. end cells, 440 cells and 251 foci analysed. (D) Sample fluorescence microscopy images of low level DprA-GFP foci within cells of R4415 (*comC0, CEP_lac_-dprA^QNQ^-gfp, dprA::spc*) and R4429 (*comC0, CEP_lac_-dprA-gfp, dprA::spc, recA::cat)* strains 15 minutes after competence induction and 5 minutes after DNA addition (250 ng µL^-1^). Scale bars, 1 µm. (E) Low level DprA-GFP foci change localisation profile in the absence of *recA* of in a *dprA^QNQ^-gfp* mutant which cannot interact with RecA. Representations focus density maps as described in Figure 1C. R4415, 7,912 cells and 6,723 foci analysed, R4429, 35,318 cells and 10,974 foci analysed. (F) Foci localisation on density heat maps as described in *panel C* for R4415 (*comC0, CEP_lac_-dprA^QNQ^-gfp, dprA::spc*) and R4429 (*comC0, CEP_lac_-dprA-gfp, dprA::spc, recA::cat*) strains. Data used as in *panel E*. Microscopy images represent sample images of each cell category showing preferential focus localisation. Scale bars, 1 µm. R4415: Small cells, 2,729 cells and 1,233 foci analysed; medium cells, 2,180 cells and 1,809 foci analysed; large cells, 2,132 cells and 2,461 foci analysed; cons. start cells, 131 cells and 135 foci analysed; cons. middle cells, 559 cells and 768 foci analysed; cons. end cells, 181 cells and 317 foci analysed. R4429: Small cells, 14,338 cells and 2,988 foci analysed; medium cells, 9,534 cells and 3,294 foci analysed; large cells, 7,508 cells and 2,925 foci analysed; cons. start cells, 727 cells and 270 foci analysed; cons. middle cells, 2,534 cells and 1,079 foci analysed; cons. end cells, 740 cells and 418 foci analysed.

### DprA anchoring at midcell is dependent on RecA

To gain further insight into the formation of tDNA-dependent DprA-GFP foci at midcell in competent cells, we reproduced these localisation experiments in strains disrupted in three genes involved in different stages of the transformation process, *i.e*. *comEC*, *ssbB* and *radA*. ComEC is proposed to form a channel in the cell membrane enabling ssDNA transfer into the cytoplasm^41^. Only 2 % of *comEC^-^*cells exhibited tDNA-dependent DprA-GFP foci, demonstrating that assembly of these foci depends on tDNA internalisation (Extended Figure 2D). Of note, it can be inferred from this result that DprA-GFP foci in competent cells grown without tDNA (Figure 2) result from the internalisation of DNA released in the medium from lysed cells. SsbB contributes to the protection and storage of internalised ssDNA to foster multiple chromosomal recombination events^42, 43^, and RadA is a helicase that extends ssDNA integration at RecA-directed D-loop intermediates^26^. Despite these key roles in transformation, neither was found to be involved in midcell DprA-GFP foci formation (Extended Figure 3).

**Figure 3:**
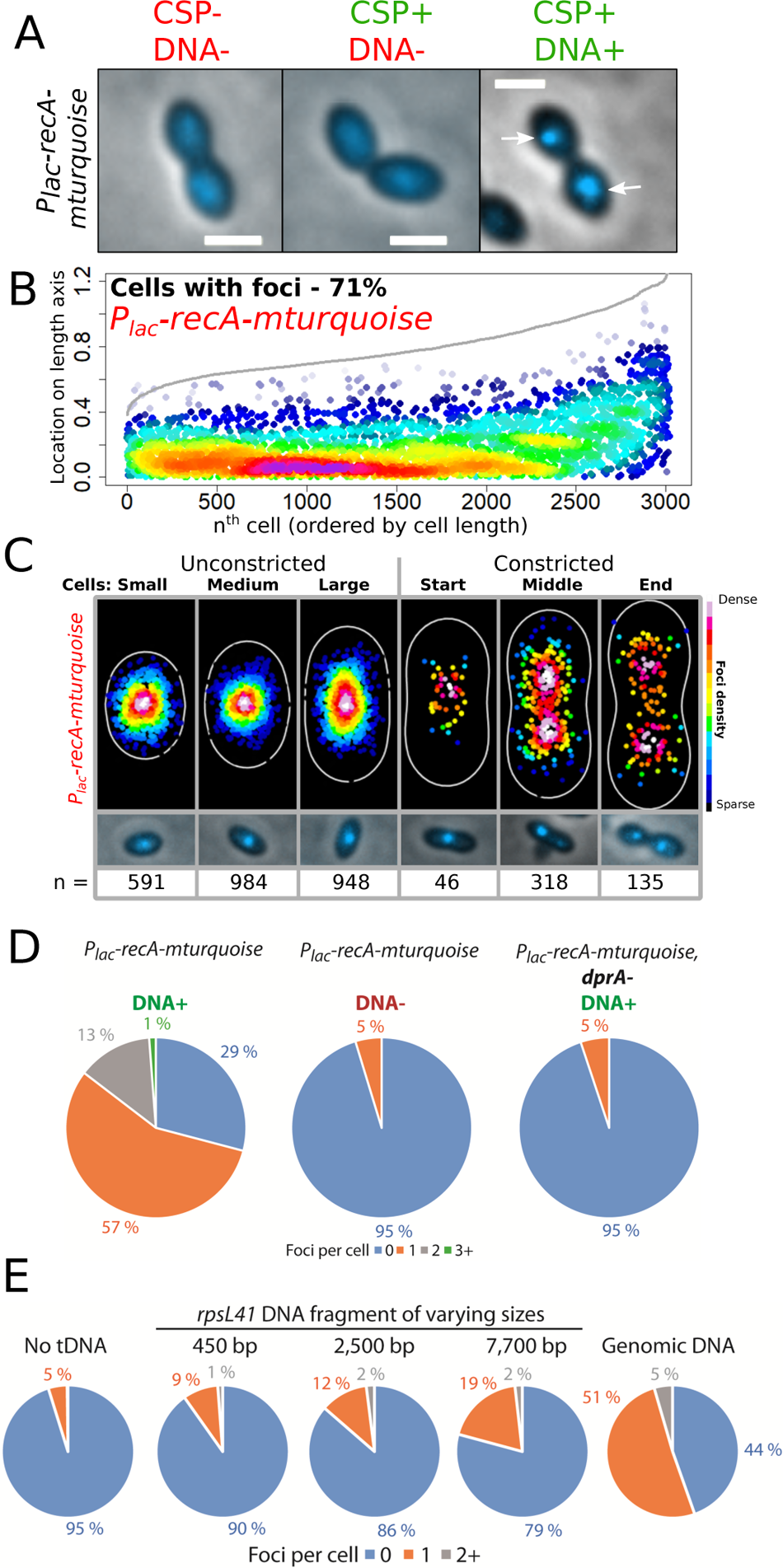
Mixed filaments of RecA/RecA-mTurquoise accumulate at midcell during competence, dependent on tDNA. (A) Sample microscopy images of RecA/RecA-mTurquoise mixed filaments in non-competent cells and competent cells in presence or absence of homologous tDNA. Images taken 15 minutes after competence induction. Strain used, R4848 (*comC0, P_lac_-recA-mturquoise*). White arrows, midcell RecA-mTurquoise foci. (B) Mixed filaments of RecA/RecA-mTurquoise accumulate at midcell, reminiscent of DprA. Representations as focus density maps as described in Figure 1C. *P_lac_-recA-mturquoise*, 3,481 cells and 3,022 foci analysed. (C) RecA/RecA-mTurquoise mixed filaments accumulate at midcell in transforming cells. Representations on density heat maps as described in Figure 2C. Microscopy images represent sample images of each cell category showing preferential focus localisation. Small cells, 923 cells and 591 foci analysed; medium cells, 1,242 cells and 984 foci analysed; large cells, 958 cells and 948 foci analysed; cons. start cells, 45 cells and 46 foci analysed; cons. middle cells, 229 cells and 318 foci analysed; cons. end cells, 84 cells and 135 foci analysed. (D) Formation of RecA-mTurquoise foci expressed from *P_lac_-recA-mturquoise* ectopic expression platform in merodiploid cells is dependent on transforming DNA and DprA. Strains used, *P_lac_-recA-mturquoise*, R4848 (*comC0, CEP_lac_-recA-mturquoise*); *P_lac_-recA- mturquoise*, *dprA^-^*, R4851, (*comC0, CEP_lac_-recA-mturquoise, dprA::spc*). (E) Formation of RecA- mTurquoise foci expressed from *P_lac_-recA-mturquoise* ectopic expression platform in merodiploid cells varies depending on the length of tDNA fragments used. Strain used, *P_lac_- recA-mturquoise*, R4848 (*comC0, CEP_lac_-recA-mturquoise*).

Next, we further explored DprA-GFP localisation with two DprA point mutants, both severely impaired in transformation and differentially altered in DprA properties: DprA^AR^, defective in dimerisation and cooperative interaction with ssDNA and DprA^QNQ^, disrupted in RecA interaction^24^. Only 2 % of transforming cells expressing the DprA^AR^-GFP fusion from the ectopic *P_lac_-dprA^AR^-gfp* construct possessed foci (Extended Figure 2D). In contrast, the DprA^QNQ^-GFP fusion still formed tDNA-dependent midcell foci. However, these were observed in fewer cells than in an isogenic wildtype DprA-GFP fusion and their localisation appeared markedly different (Figure 2DE and Extended Figure 2E). This difference can be clearly seen when the data is represented as heatmaps, with cells split into six size categories. DprA^QNQ^- GFP accumulated at the extremities of the lateral cellular axis in small and medium sized cells, and at cell poles in large cells or at the constriction site and/or at the pole in constricted cells (Extended Figure 2F). Thus, the localisation patterns of DprA-GFP and DprA^QNQ^-GFP foci markedly differ: the later appear to be excluded from the cellular areas where the former form. This result strongly suggested that DprA interaction with RecA is key for the tDNA- dependent midcell accumulation of DprA-GFP. To confirm this, we repeated the experiment with the wildtype DprA-GFP fusion in a *recA^-^* mutant, and results were similar to those observed with DprA^QNQ^-GFP (Figure 2DEF and Extended Figure 2E). These results show that the accumulation of DprA-GFP at midcell depends on DprA interaction with RecA and tDNA. Thus, this midcell accumulation point appears to attract a trio of cross-interacting partners, i.e. DprA, RecA and tDNA. Importantly, when RecA or DprA are absent, most internalised ssDNA molecules are rapidly degraded^30^. Transforming cells of a *recA comEC* double mutant showed almost no DprA-GFP foci (Extended Figure 2FG). This suggested that sufficient internalised ssDNA remains protected by DprA within *recA^-^* competent cells to enable DprA- GFP foci formation. Together, these results show that RecA drives midcell localisation of DprA- GFP foci.

### RecA accumulates at midcell during transformation

The dependency on RecA for the midcell localisation of DprA-GFP foci during transformation led us to analyse RecA localisation in competent cells. However, a *recA- mturquoise* fluorescent gene fusion generated at the native *recA* locus, despite being produced at wild type levels, was only partially functional in directing transformation and recombination repair of chromosomal damages (Extended Figure 4). In addition, this RecA- mTurquoise fusion accumulated at the cell poles during competence and generated DprA foci at this cell location when *dprA-yfp* was expressed at low concentrations, which were not formed in a RecA^+^ background (Extended Figure 5). In an attempt to visualise RecA localisation under fully functional recombination conditions, we placed the *recA-mturquoise* construct under the control of IPTG-inducible P*_lac_* promoter at the ectopic chromosomal CEP locus, allowing the production of a mixture of RecA and RecA-mTurquoise proteins in the cells, a strategy successfully used in various species^44–46^. This merodiploid strain, referred to as *recA*/*recA-mturquoise*, was equally as proficient in transformation and genome maintenance as the wildtype strain (Extended Figure 4). In this context, we found that RecA-mTurquoise accumulates into fluorescent foci at midcell in competent cells, dependent on tDNA, displaying the same foci localisation profile as cells expressing low-level DprA-GFP in the same conditions (Figure 3ABC). This result strongly suggests that midcell foci represented a functional cellular localisation for RecA and DprA during transformation. Indeed, formation of RecA-mTurquoise midcell foci was found to be dependent on tDNA and DprA (Figure 3D). Repeating this experiment in an IPTG gradient revealed that reducing the cellular levels of RecA-mTurquoise reduced the number of transformed competent cells with foci (Extended Figure 6). In addition, reducing the length of tDNA fragment reduced the number of cells presenting RecA-mTurquoise foci (Figure 3E). This interdependency between DprA and RecA for their midcell accumulation highlights the role of DprA in HR as a mediator of RecA loading on ssDNA at this precise cell location. Finally, we also attempted to directly visualise internalised ssDNA by fluorescent labelling as previously described in *B. subtilis*^47^. However, we were unable to conclusively visualise fluorescently labelled tDNA internalised in pneumococcal competent cells, which was essentially found randomly retained into multiple patches on the cell surface or in the periplasmic space as reported recently with *B. subtilis*^19^ (See Supplementary Results and Extended Figure 7). Together, these localisation studies of DprA and RecA in transforming competent cells revealed their interdependent assembly into foci at midcell, which are functionally linked to their concerted role in directing the early HR steps of transformation.

**Figure 4:**
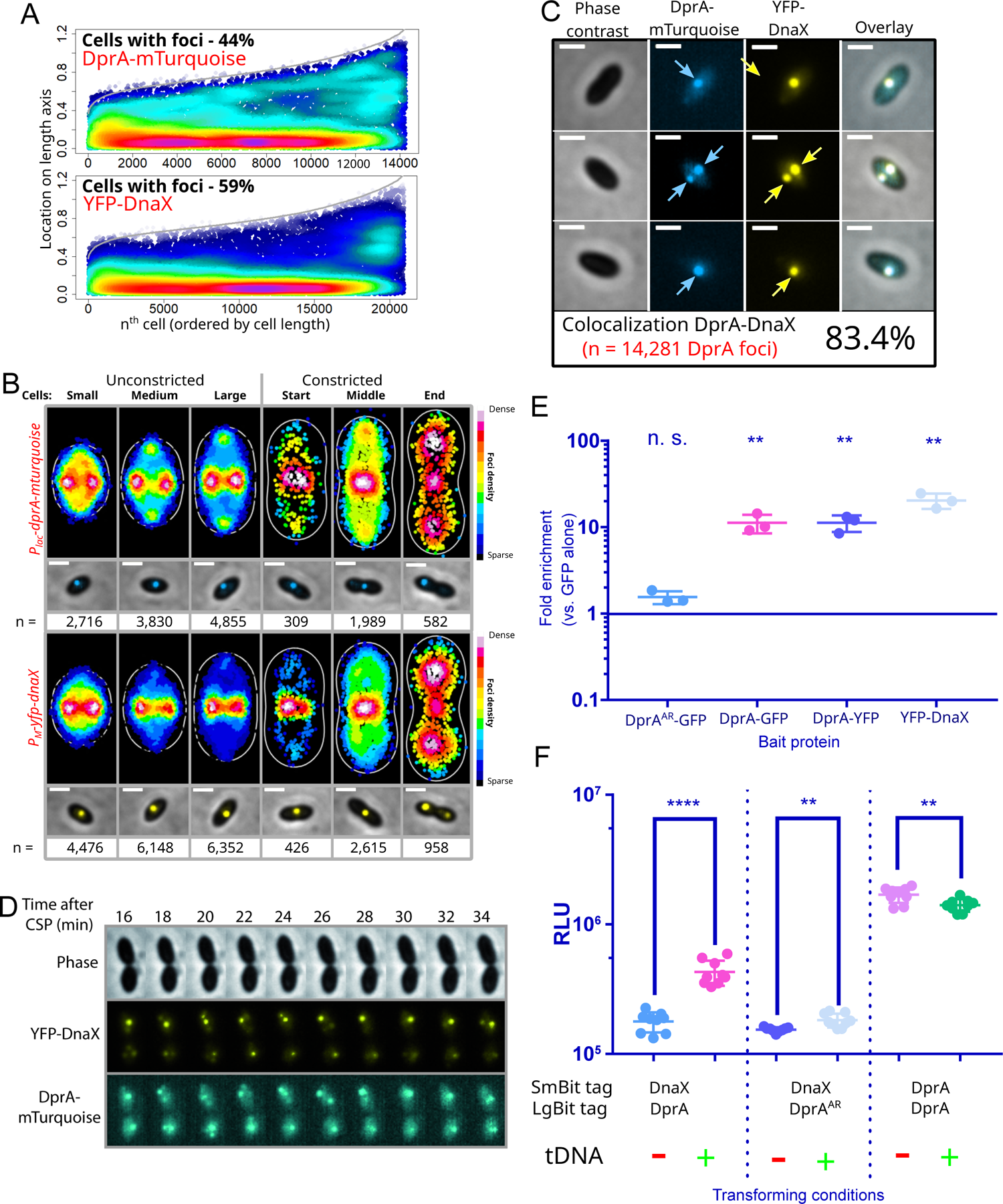
HR intermediates of transformation interact with active chromosomal replication forks in competent cells. (A) Low level DprA-mTurquoise foci display a localisation profile similar to a YFP-DnaX fluorescent fusion of the replisome clamp loader expressed in the same R4631 cells (*comC0, CEP_M_-yfp-dnaX, CEPIIP_lac_-dprA-mTurquoise, dprA^-^*). Representations as focus density maps as described in Figure 1C. 29,942 cells analysed possessing 14,281 DprA- mTurquoise foci and 20,975 YFP-DnaX foci analysed. (B) DprA-mTurquoise and YFP-DnaX foci localisation on density heat maps as described in Figure 2C for R4631 strain. Data used as in *panel A*. Microscopy images represent sample images of each cell category showing preferential focus localisation. Small cells, 9,979 cells, 2,716 DprA-mTurquoise and 4,476 YFP- DnaX foci analysed; medium cells, 8,517 cells, 3,830 DprA-mTurquoise and 6,148 YFP-DnaX foci analysed; large cells, 7,952 cells, 4,855 DprA-mTurquoise and 6,352 YFP-DnaX foci analysed; cons. start cells, 508 cells, 309 DprA-mTurquoise and 426 YFP-DnaX foci analysed; cons. middle cells, 2,390 cells, 1,989 DprA-mTurquoise and 2,615 YFP-DnaX foci analysed; cons. end cells, 596 cells, 582 DprA-mTurquoise and 958 YFP-DnaX foci analysed. (C) Sample microscopy images of R4631 strain expressing low level DprA-mTurquoise and YFP-DnaX and colocalisation of DprA-mTurquoise with YFP-DnaX in these cells. Images taken 15 minutes after competence induction and 5 minutes after DNA addition (250 ng µL^-1^). Scale bars, 1 µm. Phase contrast, phase contrast images of cells; overlay, overlay of all 3 other images. (D) DnaX and DprA produce similarly dynamic foci in competent, transforming cells. DprA-mTurquoise and YFP-DnaX observed during time-lapse microscopy of strain R4631 (*comC0, CEP_M_-yfp-dnaX, CEPII-P_lac_-dprA-mturquoise, dprA::spc*) starting 10 min after competence induction and five min after DNA addition. Images taken every two min. Phase, phase contrast images of cells. Overlay, a merge of the three other images. Scale bars, 1 µm. (E) The replisome clamp loader DnaX interacts with transforming ssDNA in early HR intermediates, as shown by co-purification of heterologous transforming ssDNA with YFP-DnaX at levels comparable to DprA-GFP and DprA-YFP in ChIP-PCR experiments. In contrast, DprA^AR^-GFP co-purified at levels comparable to the GFP alone negative control (with an enrichment value of 1, to which all other samples were normalised). Strains used, R2546, *comC0, CEP_X_-gfp*; R3406, *comC0, CEPM-yfp-dnaX*; R3728, *comC0, dprA-gfp*; R4046, *comC0, dprA^AR^-gfp*; R4404, *comC0, dprA-yfp*. Asterisks represent significant difference between samples (** = p < 0.005, n. s. = not significant). DprA^AR^-GFP, p = 0.52; DprA-GFP, p = 0.0029; DprA-YFP, p = 0.0019; YFP-DnaX, p = 0.0012. (F) Split-luciferase assay comparing cellular proximity of DnaX and DprA in presence or absence of tDNA. Luminescence signal increases when tDNA is added to competent cells containing *dprA-lgbit* and *dnaX-smbit* (R4856), indicating an increased proximity of these fusion proteins in the presence of tDNA. When *dprA-lgbit* is replaced by the dimerization mutant *dprA^AR^-lgbit* (R4861), the increase in luminescence upon addition of tDNA is attenuated. A strain containing *dprA-lgbit* and *P_lac_-dprA-smbit* (R4858) was used as a positive control for interaction since DprA dimerises, and shows high luminescence irrespective of tDNA addition. Each point represents an individual replicate, with 9 replicates done for each condition. RLU, relative luminescence units. Asterisks represent significant difference between samples (**** = p < 0.001, ** = p < 0.01, n. s. = not significant). *dnaX-smbit*, *dprA-lgbit,* p = 0.00009; *dnaX-smbit*, *dprA^AR^-lgbit*, p = 0.009; *CEP_lac_-dprA-smbit*, *dprA-lgbit,* p = 0.0066.

**Figure 5:**
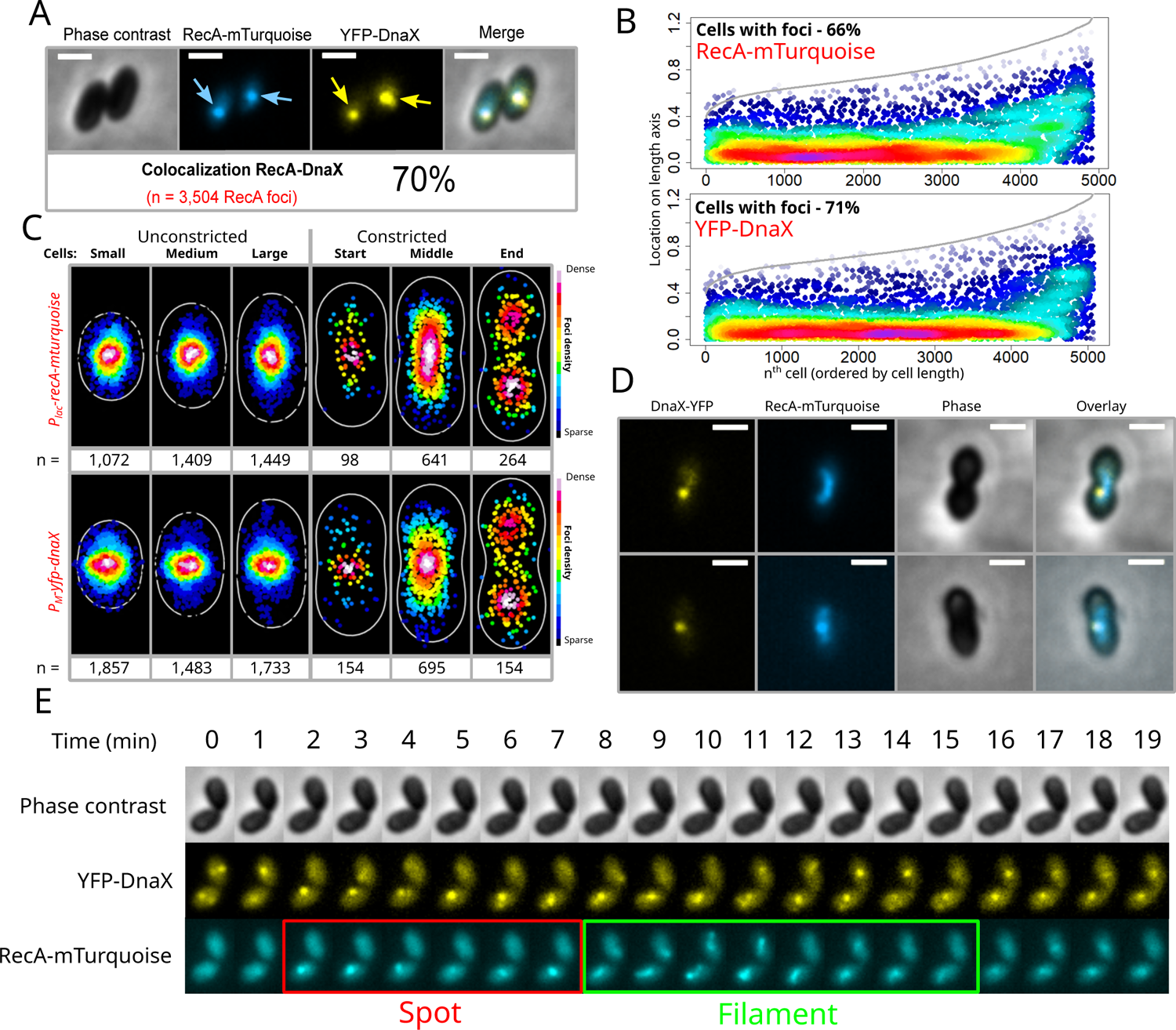
RecA/RecA-mTurquoise filaments emanate from replication forks for homology search during transformation. (A) RecA-mTurquoise and YFP-DnaX colocalise in competent cells in the presence of tDNA. Strain used, R4840 (*comC0, ssbB::luc, CEP_M_-yfp-dnaX, CEPII-P_lac_- recA-mturquoise*). 5,866 cells, 4,923 RecA-mTurquoise foci and 5,081 YFP-DnaX foci analysed. (B) Focus density maps of RecA-mTurquoise and YFP-DnaX, as described in Figure 1C. Strain, cell and foci details as in *panel A*. (C) Heatmaps of RecA-mTurquoise and YFP-DnaX as described in Figure 2C. Small cells, 1,730 cells, 1,072 RecA-mTurquoise and 1,071 YFP-DnaX foci analysed; medium cells, 1,766 cells, 1,409 RecA-mTurquoise and 1,582 YFP-DnaX foci analysed; large cells, 1,603 cells, 1,449 RecA-mTurquoise and 1,582 YFP-DnaX foci analysed; cons. start cells, 93 cells, 88 RecA-mTurquoise and 92 YFP-DnaX foci analysed; cons. middle cells, 510 cells, 1,989 RecA-mTurquoise and 614 YFP-DnaX foci analysed; cons. end cells, 165 cells, 264 RecA-mTurquoise and 257 YFP-DnaX foci analysed. (D) Microscopy images showing filaments of RecA-mTurquoise emanating from YFP-DnaX foci. (E) Time-lapse images of RecA- mTurquoise and YFP-DnaX in microfluidics experiment with time representing time after tDNA addition. Strain used as in *panel A*. dynamic extension of filaments (Figure 5E, Movie 4). In contrast to the RecA filaments formed in cells exposed to norfloxacin, those that emanate from replication forks in transformed cells were short, extending on average 0.22 (+/- 0.05) µm either side of a replisome colocalisation point. Similar short tDNA-dependent RecA filaments were observed with heterologous tDNA (genomic DNA from *E. coli*) (Extended Figure 8CDE) and, therefore, are not the result of pairing with a complementary sequence. Thus, these dynamic RecA polymers may represent presynaptic HR filaments assembled on tDNA and mediating homology search after having accessed the recipient chromosome *via* the replisome landing pad.

**Figure 6:**
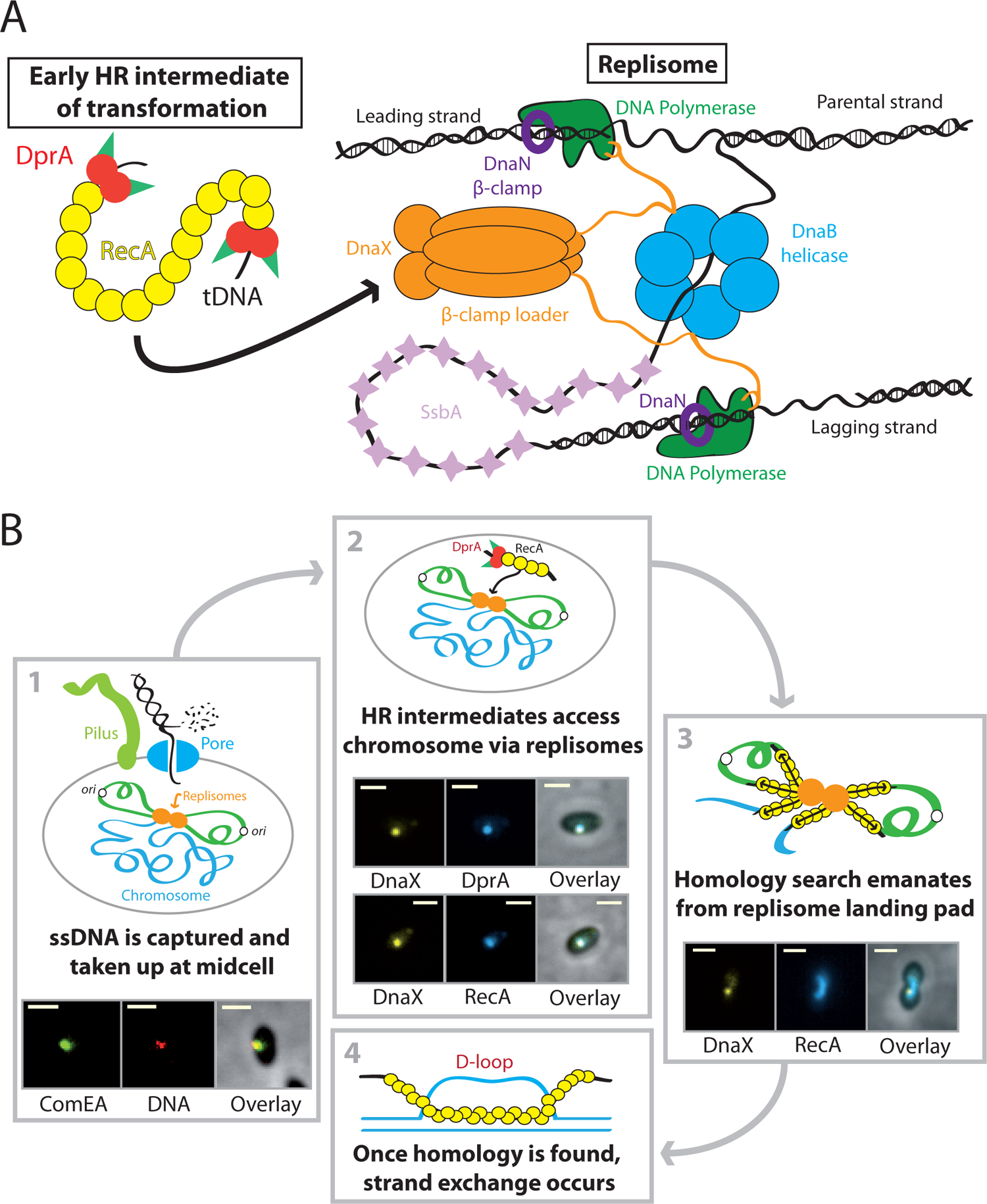
Model of the interaction between transformation and replication complexes. (A) Tight connection between early HR intermediates of transformation, comprising of the transformation-dedicated RMP DprA, the recombinase RecA and transforming ssDNA, and active replisomes during pneumococcal transformation. Diagrammatic representation of the replisome architecture based on bacterial model from *E. coli*, adapted to the pneumococcus. (B) Model of the interaction between early HR intermediates of transformation and active replisomes, and how this may facilitate homology search during HR. 1 – Transforming ssDNA is captured and internalised at midcell, providing a direct spatial link between incoming transformation nucleocomplexes and active replisomes also localised to midcell. Microscopy images taken from experiments carried out previously demonstrating colocalisation of the DNA receptor ComEA and Cy3-labelled transforming DNA at midcell^37^. Scale bar 1 µm. 2 – Early HR intermediates comprising DprA, RecA and ssDNA access the chromosome via anchoring to active replisomes. Microscopy images display the strong colocalisation of both DprA and RecA with the replisome β-clamp loader DnaX, as shown in this study. Scale bar 1 µm. 3 – RecA mediates homology search within the recipient chromosome, emanating from replisomes used as landing pads to access the chromosome. Microscopy images showing filamentation of RecA emanating from the replisome (shown via DnaX), as shown in this study. 4 – Once homology is found, strand exchange occurs between the transforming ssDNA and the homologous strand of the recipient chromosome, generating a transformation D-loop which will be resolved by replication into one wildtype and one transformed chromosome.

### DprA and RecA colocalise with chromosomal DNA replication forks in transforming cells

The localisation of midcell tDNA-dependent DprA-GFP and RecA-mTurquoise foci was very similar to the localisation of the replication forks of the chromosome tracked by using fusions to proteins of the replisome^48^. To explore whether these foci colocalised with chromosomal replication forks, a strain was generated allowing controlled, ectopic expression of both the replisomal DnaX protein fused to YFP (*CEP_M_-yfp-dnaX*; induced by maltose) and DprA-mTurquoise (*CEPII_lac_-dprA-mturquoise*; induced by IPTG). Firstly, 81 % (+/- 3,9 %) of non-competent cells possessed at least one YFP-DnaX focus, while this was slightly reduced to 73% (+/- 1,6 %) in competent cells 20 minutes after CSP addition (Extended Figure 8AB), showing that most replisomes remain intact in competent pneumococcal cells. Next, repeating the transformation experiment in this strain revealed that the cellular distribution of foci was almost identical for both fusions (Figure 4AB). 83,4 % of DprA-mTurquoise foci colocalised with YFP-DnaX foci (Figure 4C), irrespective of the cell cycle stage of the cell. The replisome protein DnaX moves dynamically around midcell^48^. Time-lapse microscopy of both DprA- mTurquoise and YFP-DnaX in competent, transforming cells showed that their midcell foci exhibited the same dynamics (Movie 1, Figure 4D). In all, this demonstrated that early DprA- mediated transformation HR intermediates navigate with the replisome. To strengthen this conclusion, ChIP-PCR experiments were carried out to explore whether YFP-DnaX was in close proximity to tDNA in transforming cells. First, results showed that a heterologous tDNA PCR fragment was copurified with DprA-GFP and DprA-YFP at 10-fold higher levels than with the DprA^AR^-GFP dimerization mutant or the unfused GFP used as a negative control (Figure 4E). Second, this tDNA was co-purified with YFP-DnaX at a similar level to DprA-GFP (Figure 4E), suggesting close proximity between early DprA-mediated HR intermediates and chromosomal replication forks during transformation. We also used a NanoBit assay^49, 50^ to explore the proximity of DprA engaged in transformation with the replisome. This system employs a luciferase separated into a large bit (LgBit) and a small bit (SmBit). Fusion of each part to different proteins that interact or are in close proximity in cells can restore luciferase activity and produce light in the presence of a furimazine-based substrate^51^. A strain possessing DprA- LgBit and an ectopic DprA-SmBit was used as a positive control and competent cells demonstrated strong luminescence irrespective of tDNA addition, due to dimerization of DprA (Figure 4F). The LgBit tag was fused to the Cter of DprA or DprA^AR^ and the SmBit was fused to the Cter of DnaX. Addition of tDNA to competent cells increased luminescence in cells coexpressing DnaX-SmBit and DprA-LgBit but not DnaX-SmBit and DprA^AR^-LgBit (Figure 4F). This result further demonstrated a close proximity between the replisome and DprA, dependent on tDNA.. Then, to formally demonstrate that RecA is also targeted to chromosomal replication forks during transformation, we analysed RecA-mTurquoise localisation in *recA*/*recA-mturquoise* competent cells co-expressing YFP-DnaX. As expected, RecA-mTurquoise foci were found to strongly colocalise with YFP-DnaX at midcell in transforming cells (Figure 5ABC).

### RecA forms dynamic tDNA-dependent filaments at chromosomal replication forks

In experiments investigating localisation of RecA-mTurquoise in the *recA*/*recA-mturquoise* strain, we observed filamentous fluorescent structures in a minority of non-competent cells (8%). These long polymers (0,82 µm +/- 0,28 µm) appear similar to RecA filaments reported as HR filaments involved in recombinational repair of double-strand breaks (DSB) in other bacteria (Badrinarayanan *et al.*, 2015; Amarh *et al.*, 2018; Wiktor *et al.*, 2021). Similarly,, exposure of non-competent pneumococcal cells to DNA damaging agent norfloxacin increased the number of cells presenting long, dynamic RecA polymers, averaging 0,84 µm (+/- 0,31 µm) in length, from 8% to 69.8% (Movies 2 and 3, Extended Figure 9). Notably, in transforming cells of this strain, we observed that 59 % (+/- 5 %) of RecA-mTurquoise foci colocalising with replisomes exhibited dynamic filaments emanating from these foci (Figure 5D). We used time-lapse microfluidics to track RecA-mTurquoise and YFP-DnaX localisation in real time in transforming cells^52^. Results showed rapid formation of RecA-mTurquoise foci in the vicinity of the replisome as little as 2 minutes after tDNA addition, and subsequent

Then, to test whether blocking DNA replication altered the capacity of DprA to mediate tDNA dependent RecA filamentation at chromosomal replication forks, we reproduced these localisation experiments in the presence of HpUra, a nucleotide analogue that selectively inhibits the essential PolC DNA polymerase of the pneumococcal replisome^30, 53^. We first analysed RecA-mTurquoise localisation in non-competent *recA*/*recA- mturquoise* cells following addition of saturating amount of HpUra that fully blocks chromosomal DNA replication and cell growth^30, 54^ (Extended Figure 10A). We observed the formation of long RecA-mTurquoise filaments (0.94 µm +/- 0.41 µm) 5 minutes after HpUra addition (Extended Figure 10B), reproducing what was observed previously in *B. subtilis*^55^. Importantly, these filaments were lost in cells lacking *recO*, showing that they depend on the RecFOR recombinase loading system^52, 53^ (Extended Figure 10B). We previously demonstrated that transformation is RecO independent^56^. Thus, we analysed RecA-mTurquoise and YFP- DnaX localisations in *recO^-^*, *recA/recAmTurquoise*, *yfp-dna*X competent cells, to prevent formation of HpUra-dependent and RecO-mediated RecA-mTurquoise filaments. Cells were exposed to HpUra for 5 minutes, then CSP was added to induce competence, and tDNA was added 10 minutes later. Cells were visualised after a further 5 minute incubation to allow tDNA internalisation. Results showed that firstly, YFP-DnaX still accumulated into midcell foci even after PolC-directed replication was blocked, showing that replisomes remained intact, although stalled (Extended Figure 10CDEF). In transforming cells with stalled replisomes, RecA still accumulated into midcell foci, which strongly colocalised with DnaX-YFP (Extended Figure 10CDEF). In conclusion, these results show that active replication is not required for RecA access to chromosomal forks during transformation and that RecO is not involved in tDNA- dependent RecA filamentation at that precise chromosomal location.

## Discussion

In this study, we reveal that the dedicated DprA-mediated and RecA-directed HR pathway of natural genetic transformation is spatiotemporally orchestrated at chromosomal replication forks in *S. pneumoniae* (Figure 6A). First, by using functional GFP fusions, we demonstrate that both DprA and RecA accumulate at midcell and colocalise with the replisome protein DnaX in a tDNA-dependent manner. These colocalisations are observed in ∼70 % of a competent, transforming population, roughly equivalent to the number of cells undergoing chromosomal replication at a given time in these growth conditions (Figures 4 and 5). Second, we found that DnaX is in physical proximity to tDNA in transforming cells (Figure 4E) and that tDNA addition promotes interaction between DnaX and DprA (Figure 4F). Interdependent DprA and RecA accumulation at chromosomal replication forks following tDNA internalisation matches the interplay between DprA, RecA and ssDNA previously uncovered by combining biochemical and genetic analyses, and proven to promote HR during transformation *via* the formation of the presynaptic HR filament^23, 24^. Strong evidence supporting this conclusion is the formation of tDNA-dependent and DprA-mediated RecA filaments emanating from the replisome (Figure 5DE). In addition, midcell DprA-GFP localisation is not observed in a *comEC* mutant (Extended Figure 2D), demonstrating the need for tDNA internalisation to generate DprA and RecA foci by interaction with the internalised ssDNA template. Third, both DprA and RecA foci and RecA filaments are observed in a minute time frame in transforming cells. These kinetics are as rapid as that of tDNA integration into the pneumococcal chromosome tracked over time with the use of a short radiolabeled tDNA fragment homologous to a specific chromosomal locus^30^. Fourth, both DprA and RecA foci and RecA filaments are formed at replication forks either with homologous or heterologous tDNA, in the same short time frame (Figure 5 and Extended Figure S8). Thus, we conclude that the RecA filaments emanating from the replication forks in transforming pneumococcal cells are presynaptic HR intermediates engaged in the search of homology on the chromosome (Figure 6B). Altogether, these findings link the HR machinery of transformation and the replisome for the first time in a naturally transformable bacterial species.

Dynamic RecA filaments formed at replication forks in transforming pneumococcal cells are similar to those of genome maintenance HR pathways visualised in single cells in several bacterial species, and demonstrated to be presynaptic filaments actively searching for homology and promoting recombinational DNA repair^44–46^. However, these RecA filaments exhibit marked differences compared to those of pneumococcal transformation reported here. Notably, the dynamic RecA filaments assembled at a single double-strand break on one copy of the neoreplicated chromosome of *Escherichia coli* or *Caulobacter crescentus* extend across the cell length, which is proposed to correspond to the bidirectional search for the uncleaved homologous DNA on the second copy of the chromosome segregated to the opposite cell pole^44–46^. We observed such long RecA filaments in growing pneumococcal cells suffering genome damages, including specific replication fork arrest caused by the HpUra PolC inhibitor as previously reported in *B. subtilis*^55^ (Extended Figure 10). In contrast, RecA filaments formed during pneumococcal transformation at replication forks are shorter (Figure 5DE). One reason for this difference may be the length of the tDNA entering the cell, evaluated to be in the range of 3 to 7 kb and gradually reduced over the competence window^43^. Another marked difference between RecA filaments in HR pathways of pneumococcal transformation and genome maintenance is their assembly site in the cell. In the latter case, RecA filaments are formed on the chromosome at the site of DNA damage, in conjunction with the formation of ssDNA template. In contrast, in the case of pneumococcal transformation, ssDNA is formed and enters the cell at the cytoplasmic membrane through ComEC, and must reach the replication fork where DprA-mediated RecA filamentation occurs. Therefore, ssDNA formation and presynaptic RecA filamentation appear to be spatially separated during transformation in the pneumococcus. Interestingly, however, tDNA capture and uptake was previously found to also occur at midcell in the pneumococcus^36, 37^. Thus, transformation appears to proceed via a midcell channel coupling DNA capture and internalisation with chromosome access and HR, which may underpin the speed at which transformation occurs in a minute time frame in the pneumococcus^30^.

Previous analysis of RecA localization during transformation in *B. subtilis* depicted a different choreography than the one reported here for *S. pneumoniae*. Interestingly, transformation occurs in non-replicating *B. subtilis* cells, as proven by the lack of DnaX-GFP foci in competent cells^57^, and GFP-RecA has been found to localise at one cell pole where the proteins directing tDNA capture and uptake accumulate^33^. Upon transformation, GFP-RecA has been found to generate long filaments from the cell pole, proposed to represent homology search on chromosomal DNA^33^. However, *B. subtilis* DprA was not found to follow the same choreography as it accumulates at midcell in transforming cells^34^. These marked deviations in RecA filamentation dynamics during transformation between *S. pneumoniae* and *B. subtilis* run parallel to the difference in the timing of competence development between these two species. Pneumococcal competence is triggered in actively replicating cells in response to a large panel of stresses^35^, including genome damage^54, 58, 59^, and lasts for a short period of time of 30 minutes^60^. In contrast, competence in *B. subtilis* occurs during nutrient starvation when cells stop replicating, and lasts for several hours^8^. Thus, transformable bacterial species have evolved distinct strategies to mediate HR-mediated chromosomal integration of tDNA, depending on how competence is integrated into their cell cycle. It will be interesting to explore how other transformable species integrate the early HR steps of transformation into their varied cell cycles. The anchoring of the presynaptic HR filaments of transformation to the chromosomal replication forks of *S. pneumoniae* not only provides them immediate access to chromosomal DNA for homology search, but also to the potential actions of the large set of proteins acting at the forks, either directly in DNA replication or occasionally to repair the damaged forks. This toolbox of DNA effectors are ideally located to assist the whole HR process of transformation up to covalent linkage of tDNA to the chromosome, many steps of which remain uncharacterised. Of note, we demonstrate with HpUra-treated competent cells that the replication forks do not need to be active to act as molecular anchors for the early step of HR of transformation (Extended Figure 10). This mirrors a previous study showing that HpUra-treated competent pneumococcal cells integrate tDNA as efficiently as non-treated cells^59^. This indicates that RecA filaments spread over the genome for homology search, emanating from replication forks.

A major perspective of this study is to identify how early HR intermediates, composed of DprA and RecA bound to tDNA, are driven to the chromosomal replication forks. Many proteins are concentrated at these vital chromosomal sites, either essential or accessory to the DNA replication process. We show that RecA drives early HR intermediates to midcell (Figure 2DEF), opening up the possibility of an interaction between RecA and such a replication protein partner. One of these known accessory effectors is the RecO protein, which is known to mediate RecA loading on ssDNA gaps. However, we demonstrate RecO is not needed for replisome access of early HR intermediates or RecA filamentation at replication forks (Extended Figure 10). In addition, transformation HR effectors SsbB and RadA also played no role in this chromosome access mechanism (Extended Figure 3).

In conclusion, this study revealed that early HR intermediates of pneumococcal transformation accumulate at chromosomal replication forks. By doing so, replication forks could provide a landing pad for presynaptic filaments of HR to access the recipient chromosome and carry out homology search, optimising the speed and efficiency of transformation.

## Materials and Methods

### Bacterial strains, competence and transformation

The pneumococcal strains, primers and plasmids used in this study can be found in Table S1. Standard procedures for transformation and growth media were used^61^. In this study, cells were prevented from spontaneously developing competence by deletion of the *comC* gene (*comC0*)^62^. Pre-competent cultures were prepared and transformation carried out as previously described^38^. Antibiotic concentrations (μg mL^-1^) used for the selection of *S. pneumoniae* transformants were: chloramphenicol (Cm), 4.5; erythromycin, 0.05; kanamycin (Kan), 250; spectinomycin (Spc), 100; streptomycin (Sm), 200; trimethoprim (Trim), 20. GraphPad Prism was used for statistical analyses. Detailed information regarding the construction of new plasmids and strains can be found in the Supplementary Information. To compare protein expression profiles, Western blots were carried out as previously described^38^. Secondary polyclonal antibodies raised against RecA and SsbB were used at 1/10,000 dilution.

### Fluorescence microscopy and image analysis

To visualise cells by epifluorescence microscopy, pneumococcal precultures grown in C+Y medium at 37 °C to an OD_550_ of 0.1 were induced with CSP (100 ng mL^-1^). Cells were incubated for 10 minutes at 37°C before addition of transforming DNA. Transforming DNA we either homologous (*S. pneumoniae* R1501 genomic DNA) or heterologous (*Escherichia coli* genomic DNA) prepared using QIAGEN 500/g Genomic tips. Cells were then incubated at 37 °C for 5 minutes unless stated. After this incubation, 2 µL samples were spotted onto a warmed microscope slide containing a slab of 1.2 % C+Y agarose as previously described ^37^ before imaging. To generate movies, images were taken of the same fields of vision at varying time points during incubation at 37 °C. Phase contrast and fluorescence microscopy was performed as previously described^63^. Images were processed using the Nis-Elements AR software (Nikon). Images were analysed using MicrobeJ, a plug-in of ImageJ^64^. Data was analysed in R and represented in two distinct ways. Firstly, focus density maps were plotted on the longitudinal axis of half cells ordered by cell length. Each spot represents the localisation of an individual focus, and spot colour represents focus density at a specific location on the half cell. Only cells with > 0 foci shown. In cells possessing > 1 foci, foci were represented adjacently on cells of the same length. Secondly, cells were separated into six categories based on cell size and presence or absence of constriction, and heatmaps were generated for each category. The six cell categories were defined in MicrobeJ to reflect those determined previously for pneumococci^63^. End of constriction (cons. end); septum = 1, circularity < 0.7; middle of constriction (cons. middle), septum = 1, 0.8 > circularity > 0.7; start of constriction (cons. start), all other cells with septum = 1; Large cells, septum = 0, cell length > 1.4 µm, circularity < 0.9; medium cells, septum = 0, 1.4 µm > cell length > 1.2 µm,0.94 > circularity < 0.9; small cells, all other cells with septum = 0. The proportions of cells found in each category were consistent with those previously observed in these conditions^63^, validating the parameters used to define the categories.

### Chromatin immunoprecipitation PCR (ChIP-PCR)

Chromatin immunoprecipitation (ChIP) was done using magnetic GFP-Trap beads as per manufacturer’s instructions (Chromotek). Briefly, cells were inoculated 1/50 in 30 mL of C+Y medium pH 7 and grown to OD_550_ 0.1. Competence was induced by addition of 100 ng mL^-1^ CSP, and cells were incubated for 10 min at 37 °C. Transforming DNA (1 kb capsule fragment absent from recipient strains amplified from D39 using primer pair DDL35-DDL36) was added at a final concentration of 1 ng µL^-1^ and cells were incubated at 37 °C for 5 minutes to allow internalisation. Cells were then fixed by addition of 3 mL Fixation solution F (50 mM Tris pH 8.0, 100 mM NaCl, 0.5 mM EGTA, 1 mM EDTA, 10 % formaldehyde) and incubation for 30 min at room temperature. Cultures were then centrifuged for 10 min at 5,000 g, 4 °C and supernantants were discarded. Pellets were washed twice in 30 mL cold PBS with centrifugation at 5,000 g, 4 °C for 10 min in between. Cells were then washed in 1 mL cold PBS and centrifuged at 10,000 g, 4 °C for 2 min before being frozen with liquid nitrogen and storage at −80 °C until use. Pellets were defrosted and resuspended in 2 mL cold Lysis L buffer (50 mM Hepes-KOH pH 7.55, 140 mM NaCl, 1 mM EDTA, 1 % triton X-100, 0.1 % Sodium deoxycholate, 100 µg mL^-1^ RNase A) before sonication in a Diagenode Bioruptor Plus sonication bath (29 cycles, 30 s sonication, 30 s rest). Resulting samples were centrifuged for 5 min at 16,000 g, 4 °C and supernatants were transferred into fresh 2 mL tubes and centrifuged for 5 min at 16,000 g, 4 °C. After transfer into fresh 2 mL tubes, 200 µL of each sample was taken and stored at −80 °C to act as a whole cell extract prior to immunoprecipitation. 25 µL of GFP-TRAP magnetic beads was then added to each sample, which was subsequently tumbled gently at 4°C for 3h 30 min. Magnetic beads were recovered by magnetism, supernatants were discarded and beads were resuspended in 1 mL cold Lysis L buffer before being centrifuged at 800 g for 5 min. Magnetic beads were recovered by magnetism, supernatants were discarded and beads were resuspended in 1 mL cold Lysis L5 buffer (50 mM Hepes-KOH pH 7.55, 500 mM NaCl, 1 mM EDTA, 1 % triton X-100, 0.1 % Sodium deoxycholate, 100 µg mL^-1^ RNase A). Magnetic beads were recovered by magnetism, supernatants were discarded and beads were resuspended in 1 mL cold Wash buffer W (10 mM Tris/HCL pH 8.0, 250 mM LiCl, 1 mM EDTA, 0.5 % NP-40, 0.5 % DOC) and beads were then recovered by magnetism and resuspended in 520 µL TES buffer (50 mM Tris/HCl pH 8.0, 10 mM EDTA, 1 % SDS). At this stage, the WCE samples were defrosted and supplemented with 300 µL TES buffer and 20 µL SDS 10 %. All samples were then incubated at 65 °C with vigorous shaking overnight. Magnetic beads were removed by magnetism and 12.5 µL proteinase K (20 mg mL^-1^) was added before incubation of the samples at 37 °C for 2 h. DNA was purified from the samples by sequential phenol:chloroform extraction and 1 µL glycogen, 40 µL sodium acetate 3M pH 5.3 and 1 mL ethanol was added before incubation at −20 °C to precipitate the DNA. Samples were then centrifuged at 16,000 g, 4 °C for 15 min and the supernatant was carefully removed before the pellets were resuspended in 100 µL TE buffer pH 8 and incubated at 65 °C for 20 min. DNA was then purified using GFX PCR purification columns (GE Healthcare). DNA samples were diluted (1/200 for WCE, 1/20 for IP samples) and probed in triplicate by qPCR for the presence of the 1 kb capsule fragment using iTaq DNA polymerase (BIO-RAD) and primer pair DDL34-DDL35, which amplify a 115 bp region within the 1 kb fragment. Specific amplifications were confirmed by single peaks in melting curve analysis. Cycle threshold (CT) values were obtained according to the software instructions. Relative quantification was performed with the 2^-ΔΔCT^ method^65^. Each PCR reaction, run in duplicate for each sample, was repeated for at least two independent times. Data are represented as mean ± s.e.m calculated from triplicate repeats, with individual data points plotted.

### Split-luciferase assay

Split luciferase assays were carried out as previously described^49, 50^, with modifications. Briefly, pneumococcal cells were grown in C+Y medium (with 50 µM IPTG where required) at 37 °C until OD_550_ 0.1 and competence was induced by addition of 100 ng mL^-1^ CSP. Cells were then incubated for 10 min at 37 °C before addition of R1501 chromosomal DNA (250 ng µL^-1^) where noted, followed by a further 5 min incubation at 37 °C. Cells were then washed in fresh C+Y medium and 1 % NanoGlo substrate (Promega) was added and luminescence was measured 20 times every 1 min in a plate reader (VarioSkan luminometer, ThermoFisher). Data are represented as mean ± s.e.m calculated from nine independent repeats, with individual data points plotted.

## Acknowledgements

We thank Isabelle Mortier-Barriere for support with microfluidics experiments. We thank Jérome Rech for support with microscopy and video assembly. We thank the LITC imaging platform of Toulouse TRI for their assistance in microscopy. This work was funded by the Centre National de la Recherche Scientifique, University Paul Sabatier and the Agence Nationale de la Recherche (grants ANR-10-BLAN-1331 and ANR-17-CE13-0031).

## Author contributions

C. J. and P. P. wrote the paper. C. J., R. H., A-L. S., M. D. and D. D. L. performed the experiments. C. J. and P. P. designed and analysed the experiments and interpreted the data.

## Extended Figure Legends

**Extended Figure 1:**
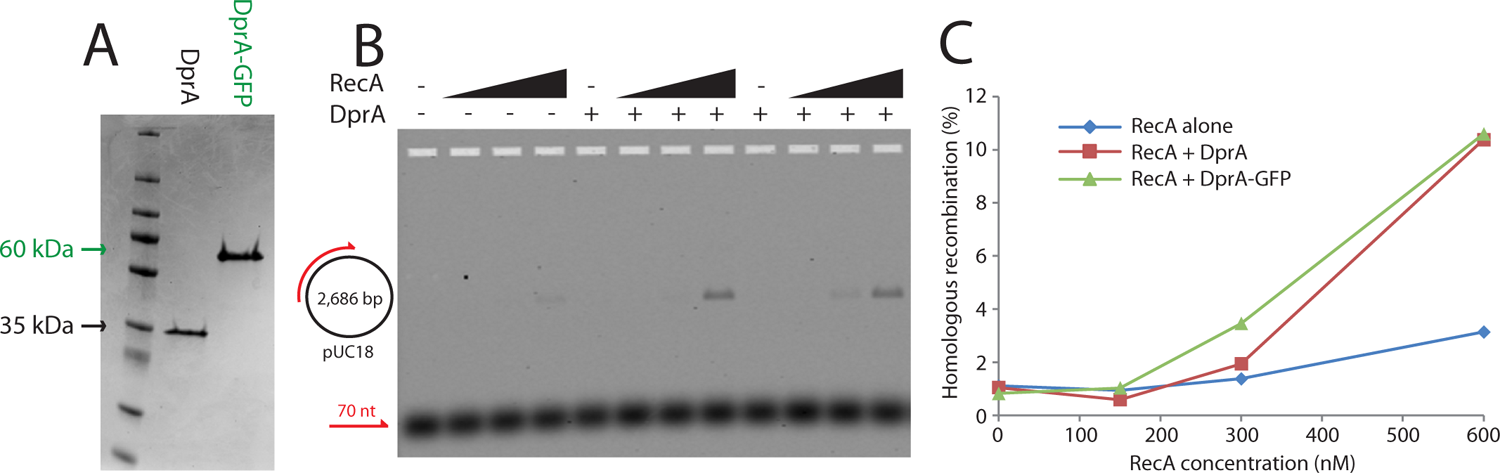
DprA-GFP is fully functional for D-loop formation in vitro. (A) Coommassie gel showing purified DprA (30 kDa) and DprA-GFP (60 kDa) proteins. (B) Purified DprA-GFP protein loads RecA onto ssDNA to stimulate D-loop formation at levels comparable to wildtype DprA. D-loop assays showing stimulation of D-loop formation between the 2,680 bp pUC18 plasmid and a Cy3-labelled 70 nucleotide (nt) fully complementary primer by varying concentrations of RecA in the presence or absence of DprA or DprA-GFP. RecA concentrations in ascending order (nM); 150, 30, 600. (C) Quantification of D-loop formation carried out on the image in *panel B*.

**Extended Figure 2:**
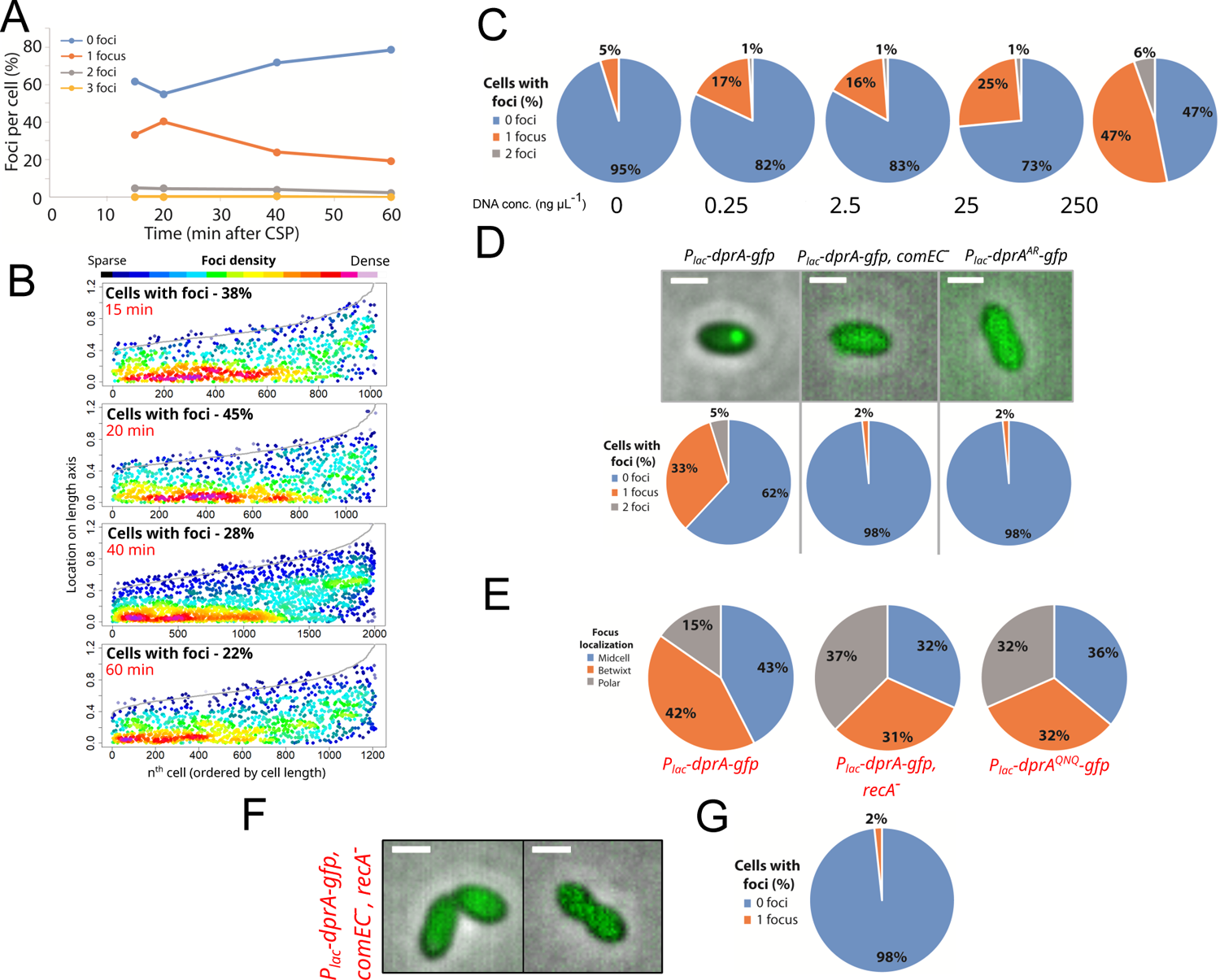
Further analysis of localisation dynamics of *P_lac_-dprA-gfp*. (A) Time-course experiment of low-level DprA-GFP foci per cell in competent cells. 15 min, 2,355 cells and 1,023 foci analysed; 20 min, 2,260 cells and 1,129 foci analysed; 40 min, 5,988 cells and 2,001 foci analysed; 60 min, 5,244 cells and 1,216 foci analysed. (B) Low level DprA-GFP foci persist at midcell up to 60 minutes after competence induction. Representations as focus density maps as described in Figure 1C. Cells and foci analysed as in *panel D*. (C) Analysis of the repartition of low-level DprA-GFP foci within competent cells in a gradient of transforming DNA. 0 ng µL^-1^ DNA, 2,621 cells and 128 foci analysed; 0.25 ng µL^-1^ DNA, 2,222 cells and 425 foci analysed; 2.5 ng µL^-1^ DNA, 6,038 cells and 1,098 foci analysed; 25 ng µL^-1^ DNA, 4,215 cells and 1,194 foci analysed; 250 ng µL^-1^ DNA, 2,190 cells and 1,313 foci analysed. (D) Sample fluorescence microscopy images and analysis of the repartition of low level DprA-GFP foci within cells of R4262 (*comC0, CEP_lac_-dprA-gfp, dprA::spc*), R4401 (*comC0, CEP_lac_-dprA-gfp, dprA::spc, comEC::ery*) and R4413 (*comC0, CEP_lac_-dprA^AR^-gfp, dprA::spc*) strains 15 minutes after competence induction and 5 minutes after DNA addition (250 ng µL^-1^). Scale bars, 1 µm. (E) Distribution of low level DprA-GFP foci in cells of R4415 (*comC0, CEP_lac_-dprA^QNQ^-gfp, dprA::spc*) and R4429 (*comC0, CEP_lac_-dprA-gfp, dprA::spc, recA::cat)* strains 15 minutes after competence induction and 5 minutes after DNA addition (250 ng µL^-1^). Data taken from Figures 2B and 2E. Midcell, betwixt and polar localisations defined as in Figure 1E. (F) Sample fluorescence microscopy image of low level DprA-GFP foci within cells of R4618 (*comC0, CEP_lac_-dprA-gfp, dprA::spc, comEC::trim, recA::cat*). Conditions as in Figure 2B. (G) Analysis of the repartition of low level DprA-GFP foci within cells of R4618. 2,455 cells and 22 foci analysed.

**Extended Figure 3:**
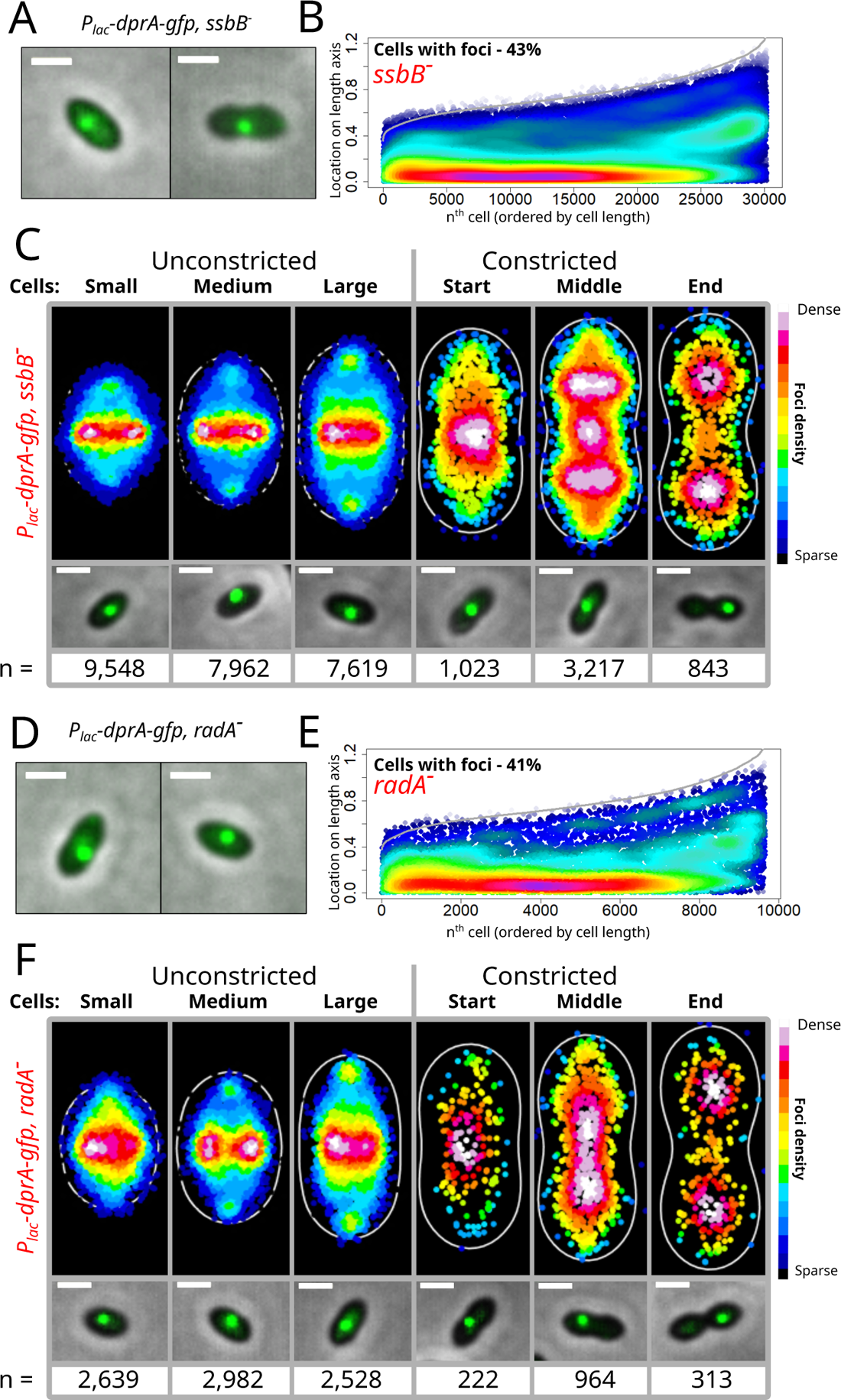
The absence of SsbB or RadA does not affect the localisation of early HR intermediates of transformation. (A) Sample fluorescence microscopy image of strain R4400 (*comC0, CEP_lac_-dprA-gfp, dprA::spc, ssbB::cat)* 15 minutes after competence induction and 5 minutes after DNA addition (250 ng µL^-1^). Scale bars, 1 µm. (B) Low cellular DprA-GFP accumulates at midcell upon addition of transforming DNA in the absence of SsbB. Representations as focus density maps as described in Figure 1C. 41,788 cells and 30,213 foci analysed. (C) DprA-GFP localisation in the absence of SsbB represented as density heat maps as in Figure 2C. Microscopy images represent sample images of each cell category showing preferential focus localisation. Scale bars, 1 µm. Small cells, 16,723 cells and 9,548 foci analysed; medium cells, 11,722 cells and 7,962 foci analysed; large cells, 9,029 cells and 7,619 foci analysed; cons. start cells, 948 cells and 1024 foci analysed; cons. middle cells, 2,689 cells and 3,217 foci analysed; cons. end cells, 677 cells and 843 foci analysed. (D) Sample fluorescence microscopy image of strain R4625 (*comC0, CEP_lac_-dprA-gfp, dprA::spc, radA::trim)* 15 minutes after competence induction and 5 minutes after DNA addition (250 ng µL^-1^). Scale bars, 1 µm. (E) Low cellular DprA-GFP accumulates at midcell upon addition of transforming DNA in the absence of RadA. Representations as described in Figure 1C. 17,925 cells and 9,667 foci analysed. (F) DprA-GFP localisation in the absence of RadA represented as density heat maps as in Figure 2C. Microscopy images represent sample images of each cell category showing preferential focus localisation. Small cells, 7,544 cells and 2,638 foci analysed; medium cells, 6,678 cells and 2,982 foci analysed; large cells, 5,315 cells and 2,528 foci analysed; cons. start cells, 421 cells and 222 foci analysed; cons. middle cells, 1,528 cells and 984 foci analysed; cons. end cells, 439 cells and 313 foci analysed.

**Extended Figure 4:**
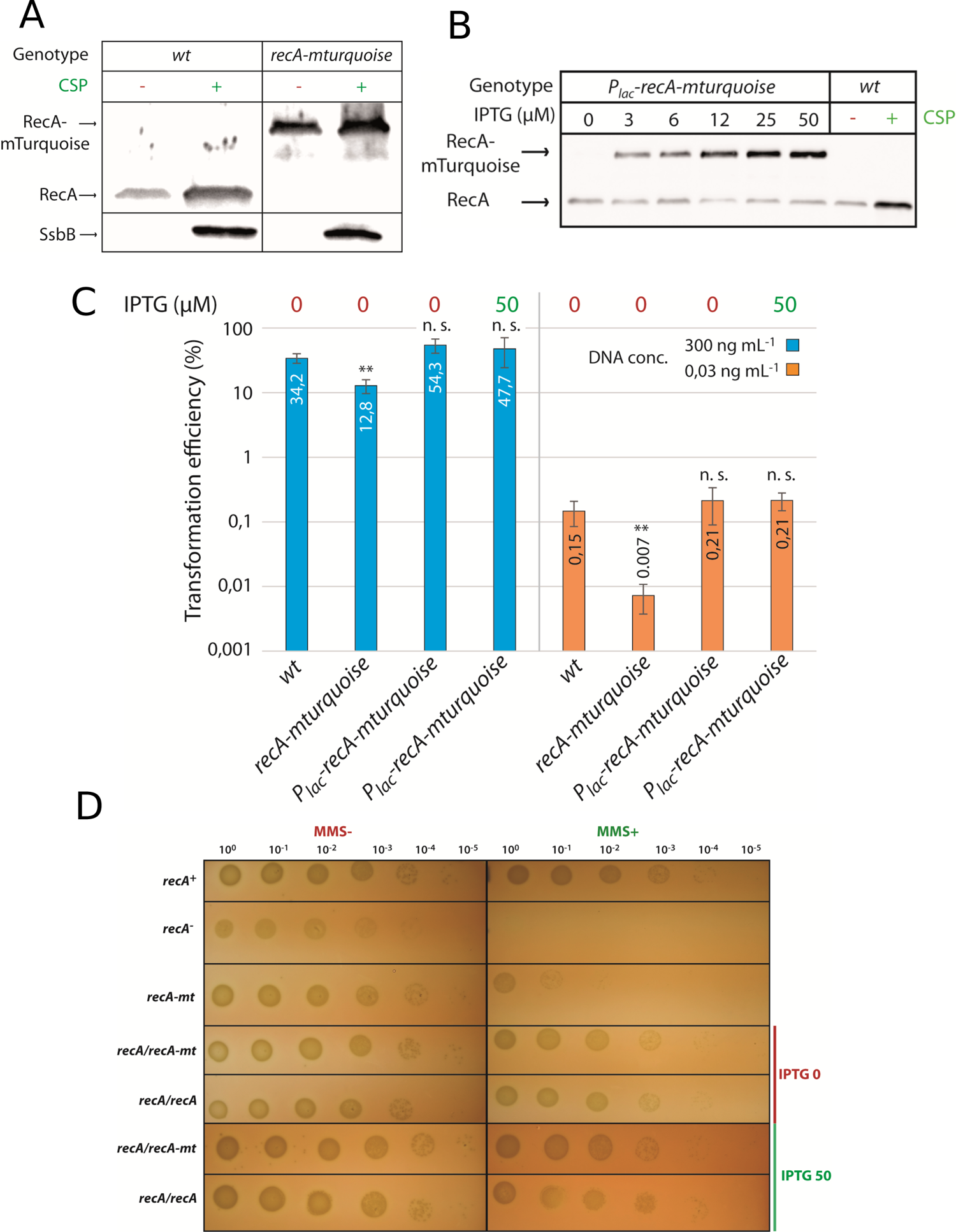
Analysis of the functionality of RecA-mTurquoise fusions. (A) Western blot probed with α-RecA antibodies showing cellular levels of RecA and RecA-mTurquoise in competent (CSP+) and non-competent (CSP-) cells. Samples also probed with α-SsbB antibodies to confirm induction of competence. Samples taken 15 minutes after competence induction. Strains used; RecA, R1501 (*comC0*); RecA-mTurquoise, R4712 (*comC0, recA- mturquoise*). (B) Western blot probed with α-RecA antibodies showing cellular levels of RecA and RecA-mTurquoise in a strain possessing *recA* and *CEP_lac_-recA-mturquoise* grown in varying concentrations of IPTG and in absence of CSP. Wildtype RecA strain included as a control in presence or absence of CSP. Strains used; RecA, R1501 (*comC0*); RecA/RecA-mTurquoise, R4848 (*comC0, CEP_lac_-recA-mturquoise*). (C) Comparison of transformation efficiencies of various *recA* mutant strains to wild type. Saturating (300 ng mL^-1^) and non-saturating (0.03 ng mL^-1^) concentrations of *rpsL41* PCR fragment, conferring streptomycin resistance via point mutation, used. DNA identity; 3,434 bp *rpsL41* PCR fragment amplified from R304 strain using MB117-MB120 primer pair. Transformation pre-cultures prepared in 0 or 50 µM IPTG as noted. Strains used: *wt*, R1501 (*comC0*); *recA-mturquoise*, R4712 (*comC0, recA-mturquoise*), *CEP_lac_-recA-mturquoise*, R4848 (*comC0, CEP_lac_-recA-mturquoise*). Data represented as mean ± s.e.m. of triplicate repeats. Asterisks represent significant difference between test samples and wildtype controls for a given DNA concentration (** = p < 0.01, n. s. = not significant). (D) Spot tests comparing growth of competent (CSP+) and non-competent (CSP-) *recA* mutants to wildtype in presence or absence of methanemethylsulfonate (MMS, 0.02 %). Strains used: *wt*, R1501 (*comC0*); *recA-*, (R4857, *comC0, recA::trim*), *recA-mturquoise*, R4712 (*comC0, recA- mturquoise*), *CEP_lac_-recA-mturquoise*, R4848 (*comC0, CEP_lac_-recA-mturquoise*); R4664 (*comC0, CEP_lac_-recA*). R4848 and R4664 grown and plated with 0 and 50 µM IPTG to compare presence and absence of *CEP_lac_-recA-mturquoise* induction.

**Extended Figure 5:**
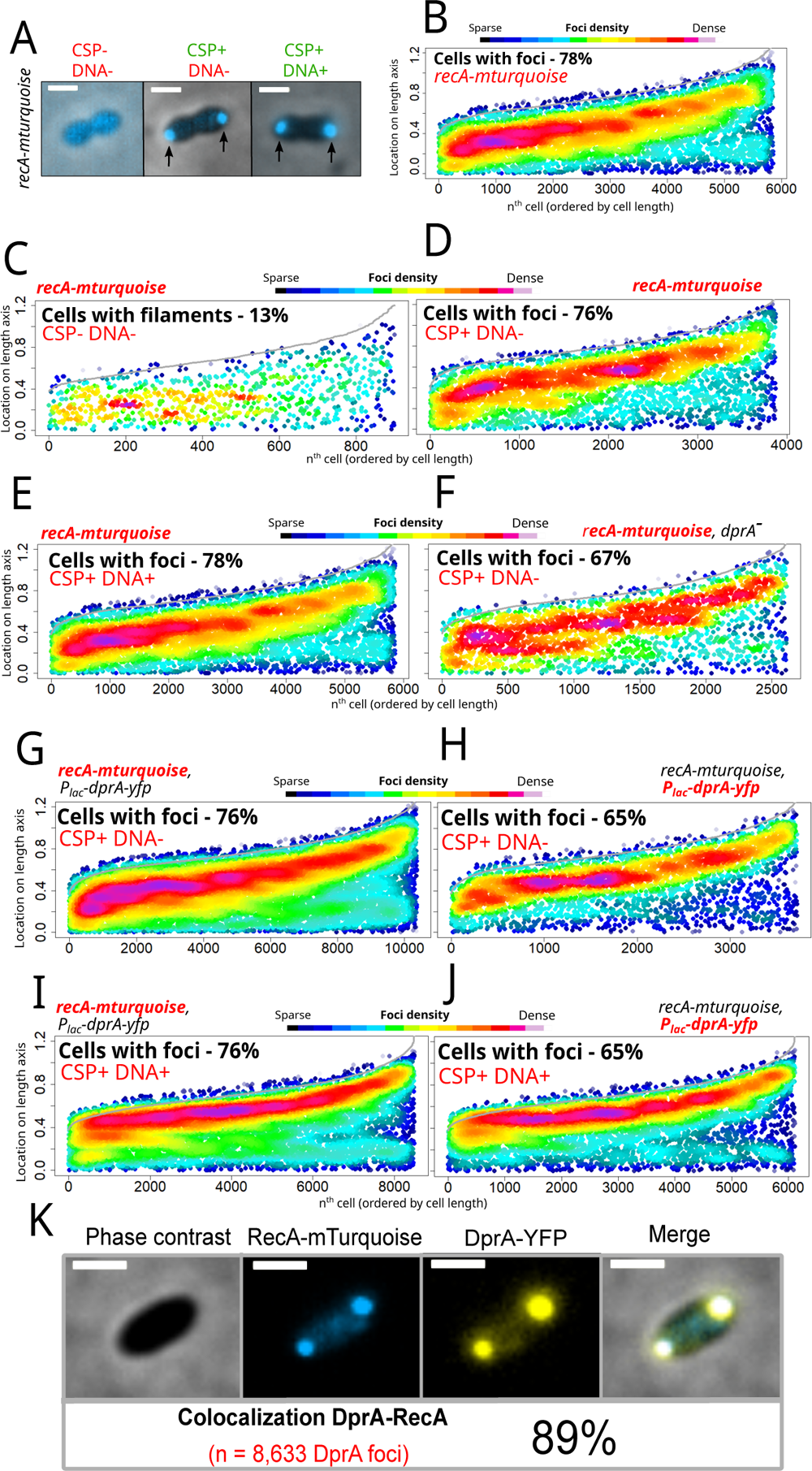
Aberrant cellular localisation of RecA-mTurquoise fusion. (A) Sample microscopy images of RecA-mTurquoise in non-competent cells and competent cells in presence or absence of homologous transforming DNA. Images taken 15 minutes after competence induction. Strain used, R4712 (*comC0, recA-mturquoise*). Black arrows, polar foci. (B) RecA-mTurquoise alone accumulates at the cell poles in a majority of transforming cells. Representations as focus density maps as described in Figure 1C. 8,125 cells and 5,858 foci analysed. (C) RecA-mTurquoise forms centrally localised bundles in a minority of non- competent cells. Representations as focus density maps as described in Figure 1C. 5,512 cells and 901 bundles analysed. (D) RecA-mTurquoise forms foci at the cell poles in competent cells in absence of transforming DNA. Representations as focus density maps as described in Figure 1C. 6,066 cells and 3,874 foci analysed. (E) The presence of transforming DNA does not alter the polar localisation of RecA-mTurquoise. Representations as focus density maps as described in Figure 1C. 8,125 cells and 5,858 foci analysed. (F) The absence of DprA does not alter RecA-mTurquoise localisation in competent cells. Representations as focus density maps as described in Figure 1C. 6,066 cells and 3,874 foci analysed. (G) RecA-mTurquoise forms polar foci in a strain expressing DprA-YFP in the presence of transforming DNA. Representations as focus density maps as described in Figure 1C. 9,337 cells, 10,367 RecA- mTurquoise foci analysed. Strain used, R4742, *comC0*, *P_lac_-dprA-yfp*, *recA-mturquoise*, *dprA*::*spc*. (H) DprA-YFP forms polar foci in a strain expressing RecA-mTurquoise in the presence of transforming DNA. Strain and images used as in *panel E*. Representations as focus density maps as described in Figure 1C. 9,337 cells, 3,721 DprA-YFP foci analysed. (I) RecA- mTurquoise forms polar foci in a strain expressing DprA-YFP in the presence of transforming DNA. Representations as focus density maps as described in Figure 1C. 7,074 cells, 8,493 RecA-mTurquoise foci analysed. Strain used as in *panel E*. (J) DprA-YFP forms polar foci in a strain expressing RecA-mTurquoise in the presence of transforming DNA. Images used as in *panel G*. Representations as focus density maps as described in Figure 1C. 7,074 cells, 6,120 DprA-YFP foci analysed. Strain used as in *panel E*. (K) Sample microscopy images of a strain expressing low level DprA-YFP and RecA-mTurquoise in competent, transforming cells and colocalisation of DprA-YFP with RecA-mTurquoise in these cells. Images taken 15 minutes after competence induction and 5 minutes after DNA addition (250 ng µL^-1^). Scale bars, 1 µm. Phase, phase contrast images of cells; overlay, overlay of all 3 other images. Strain used as in *panels I* and *J*.

**Extended Figure 6:**
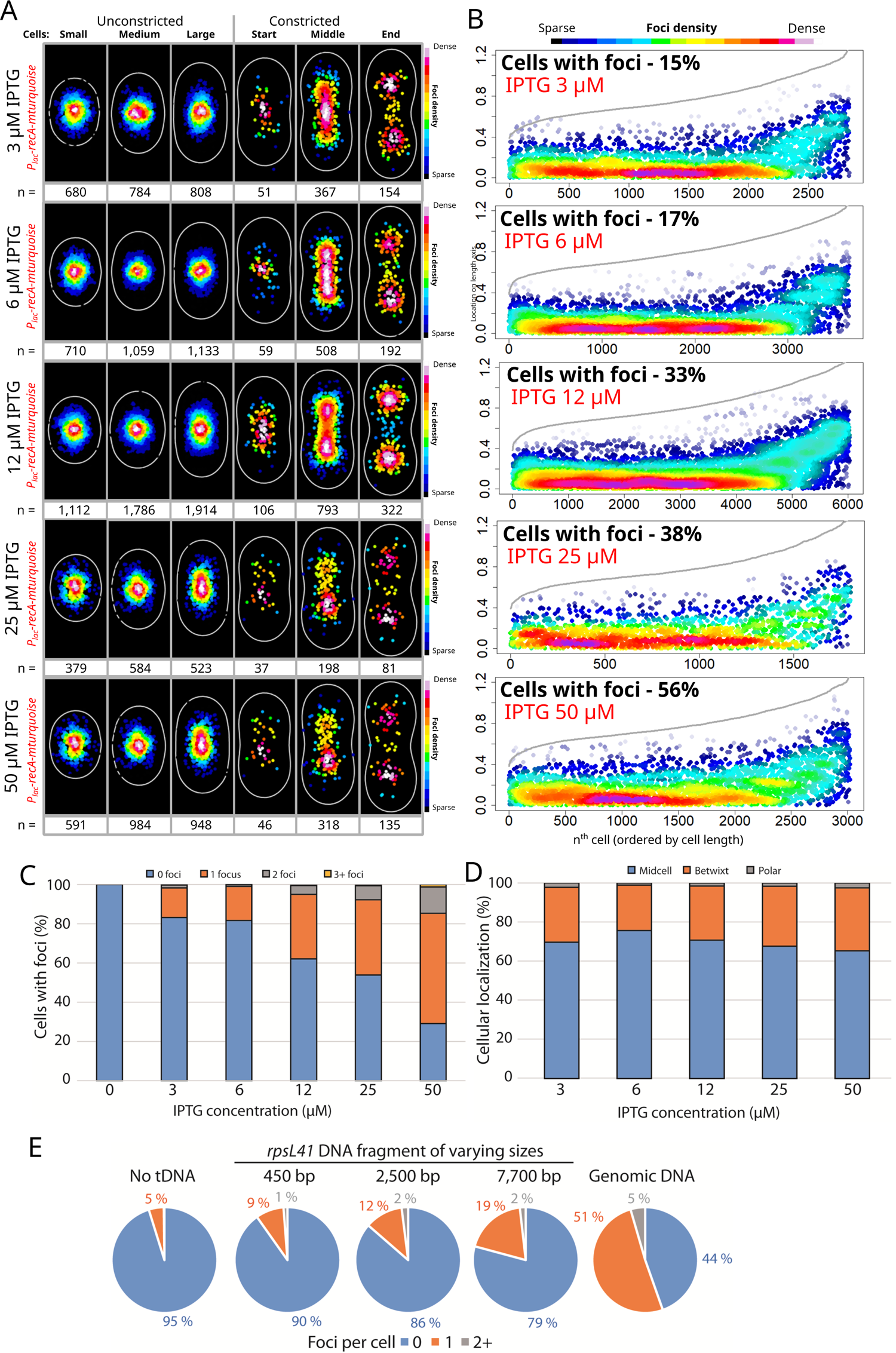
Reducing cellular levels of RecA-mTurquoise in a mixed filament strain reduces midcell accumulation in transforming cells. (A) Heatmaps of RecA-mTurquoise foci in competent, transforming RecA/RecA-mTurquoise cells grown in varying IPTG concentrations of IPTG (3-50 µM), as described in Figure 2C. Strain used: R4848, *comC0, CEP_lac_-recA-mturquoise*. 3 µM IPTG, Small cells, 7,056 cells and 680 foci analysed; medium cells, 3,594 cells and 784 foci analysed; large cells, 3,012 cells and 808 foci analysed; cons. start cells, 282 cells and 51 foci analysed; cons. middle cells, 964 cells and 367 foci analysed; cons. end cells, 321 cells and 154 foci analysed. 6 µM IPTG, Small cells, 8,533 cells and 710 foci analysed; medium cells, 4,617 cells and 1,059 foci analysed; large cells, 3,911 cells and 1,133 foci analysed; cons. start cells, 321 cells and 59 foci analysed; cons. middle cells, 1,238 cells and 508 foci analysed; cons. end cells, 362 cells and 192 foci analysed. 12 µM IPTG, Small cells, 6,037 cells and 1,112 foci analysed; medium cells, 3,424 cells and 1,786 foci analysed; large cells, 3,089 cells and 1,914 foci analysed; cons. start cells, 229 cells and 106 foci analysed; cons. middle cells, 854 cells and 793 foci analysed; cons. end cells, 271 cells and 322 foci analysed. 25 µM IPTG, Small cells, 1,187 cells and 379 foci analysed; medium cells, 1,002 cells and 584 foci analysed; large cells, 806 cells and 523 foci analysed; cons. start cells, 49 cells and 37 foci analysed; cons. middle cells, 186 cells and 198 foci analysed; cons. end cells, 66 cells and 81 foci analysed. 50 µM IPTG, Small cells, 923 cells and 591 foci analysed; medium cells, 1,242 cells and 984 foci analysed; large cells, 958 cells and 948 foci analysed; cons. start cells, 45 cells and 46 foci analysed; cons. middle cells, 229 cells and 318 foci analysed; cons. end cells, 84 cells and 135 foci analysed. (B) Focus density maps of RecA-mTurquoise foci in competent, transforming RecA/RecA-mTurquoise cells as described in Figure 2B. Strains, conditions and images used as in *panel A*. 3 µM IPTG, 15,229 cells and 2,844 foci analysed; 6 µM IPTG, 18,982 cells and 3,661 foci analysed; 12 µM IPTG, 13,940 cells and 6,033 foci analysed; 25 µM IPTG, 3,296 cells and 1,802 foci analysed; 50 µM IPTG, 3,481 cells and 3,022 foci analysed. (C) Percentage of RecA-mTurquoise foci in RecA/RecA-mTurquoise cells grown in varying IPTG concentrations. Strains, conditions and images used as in *panel A*. (D) Cellular localisation of RecA-mTurquoise foci in RecA/RecA-mTurquoise cells grown in varying IPTG concentrations. Strains, conditions and images used as in *panel A*. (E) RecA-mTurquoise foci per cell in transformation experiments using DNA fragments of varying sizes. Strain used, R4848 (*comC0, CEP_lac_-recA-mturquoise*).

**Extended Figure 7:**
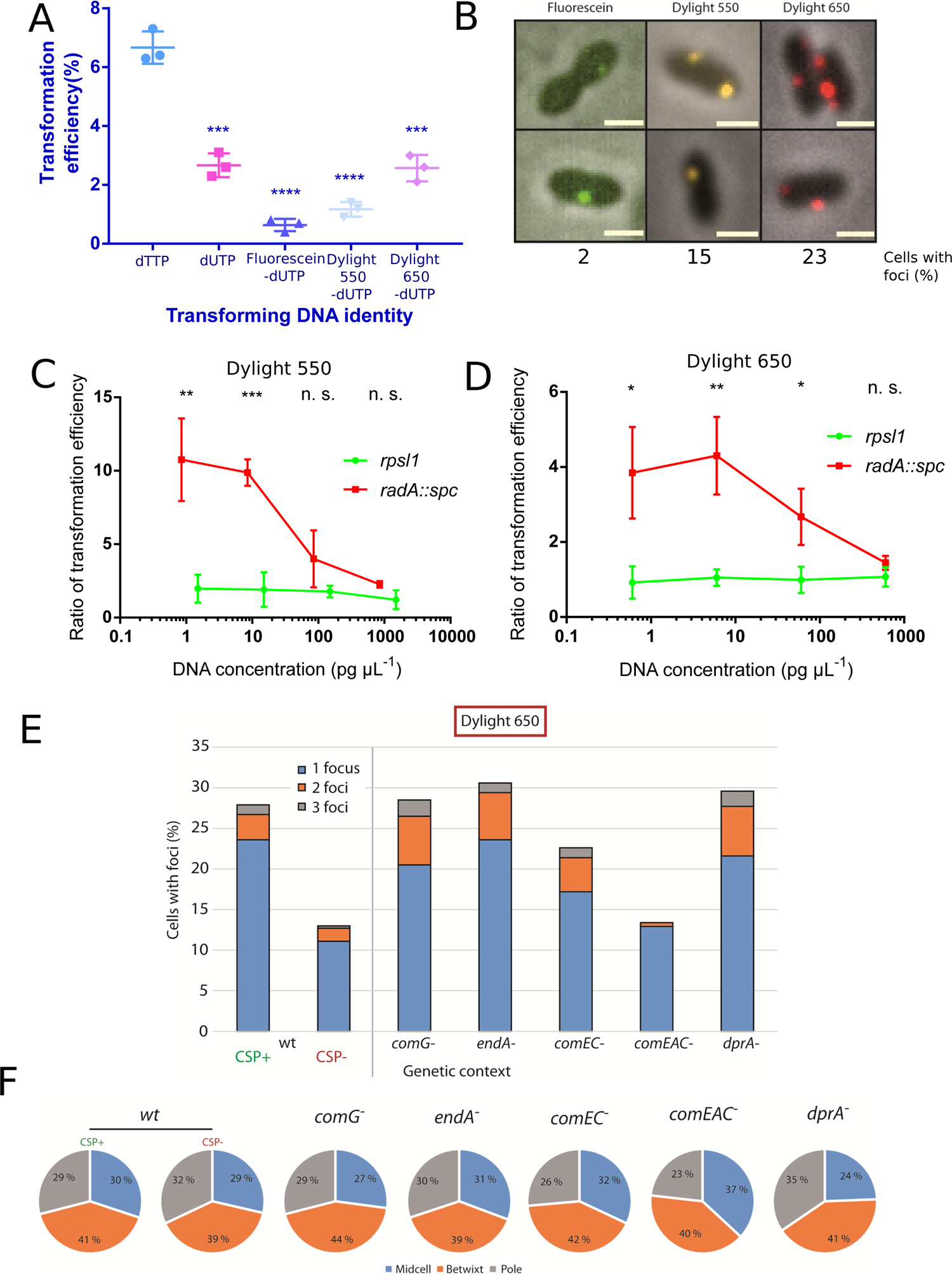
Exploration of fluorescent DNA internalisation during pneumococcal transformation. (A) Comparison of transformation of *rpsL41* PCR fragments containing either dTTP or dUTP. dUTP bases were either unlabelled, labelled with fluorescein, or labelled with amino-allyl and tagged with Dylight 550 or 650 fluorophores. Asterisks represent significant difference between test samples and dTTP control (*** = p < 0.005, **** = p < 0.001). dUTP, p = 0.0005; d-UTP-fluorescein, p < 0.0001; d-UTP-Dylight 550, p < 0.0001; d-UTP-Dylight 650, p = 0.0006. (B) Comparison of microscopy images of competence cells of R1501 (*comC0*) transformed with DNA labelled with fluorescein, Dylight 550 or 650. Analysis of microscopy images of cells transformed with DNA labelled with various fluorophores, showing cells possessing detectable fluorescent foci. (C) Comparison of transformation efficiency of Dylight 550 transformed with PCR fragments of *rpsL41* or *radA::spc* in a concentration gradient of transforming DNA. Values plotted represent ratios of transformation efficiency between labelled and labelled PCR fragments. Strain used as in *panel A.* Asterisks represent significant difference ratios for each PCR fragment (** = p < 0.01, *** = p < 0.005, n. s., not significant). ∼1 pg µL^-1^ DNA, p = 0.0069; ∼10 pg µL^-1^ DNA, p = 0.0007; ∼100 pg µL^-1^ DNA, p = 0.12; ∼1,000 pg µL^-1^ DNA, p = 0.056. (D) Comparison of transformation efficiency of Dylight 650 transformed with PCR fragments of *rpsL41* or *radA::spc* in a concentration gradient of transforming DNA. Values plotted as in *panel C*. Strain used as in *panel A.* Asterisks represent significant difference ratios for each PCR fragment (* = p < 0.05, ** = p < 0.01, n. s., not significant). ∼1 pg µL^-1^ DNA, p = 0.017; ∼10 pg µL^-1^ DNA, p = 0.006; ∼100 pg µL^-1^ DNA, p = 0.025; ∼1,000 pg µL^-1^ DNA, p = 0.11. (E) Comparison of foci present in cells of various transformasome mutants after exposure to Dylight 650-labelled *rpsL41* PCR fragments. Strains used: wt, R1501 (*comC0*); *comG^-^*, R4655 (*comC0, comC-luc, comG::kan*); *endA^-^*, R2811 (*comC0, endA::cat*); *comEC^-^*, R2586 (*comC0, comEC::ery*); *comEAC^-^*, R4653 (*comC0, comC-luc, comEAC::spc*); *dprA^-^*, R2018 (*comC0, dprA::spc*). (F) Comparison of focus localisation in cells of various transformasome mutants after exposure to Dylight 650-labelled *rpsL41* PCR fragments. Strains used as in *panel E*.

**Extended Figure 8:**
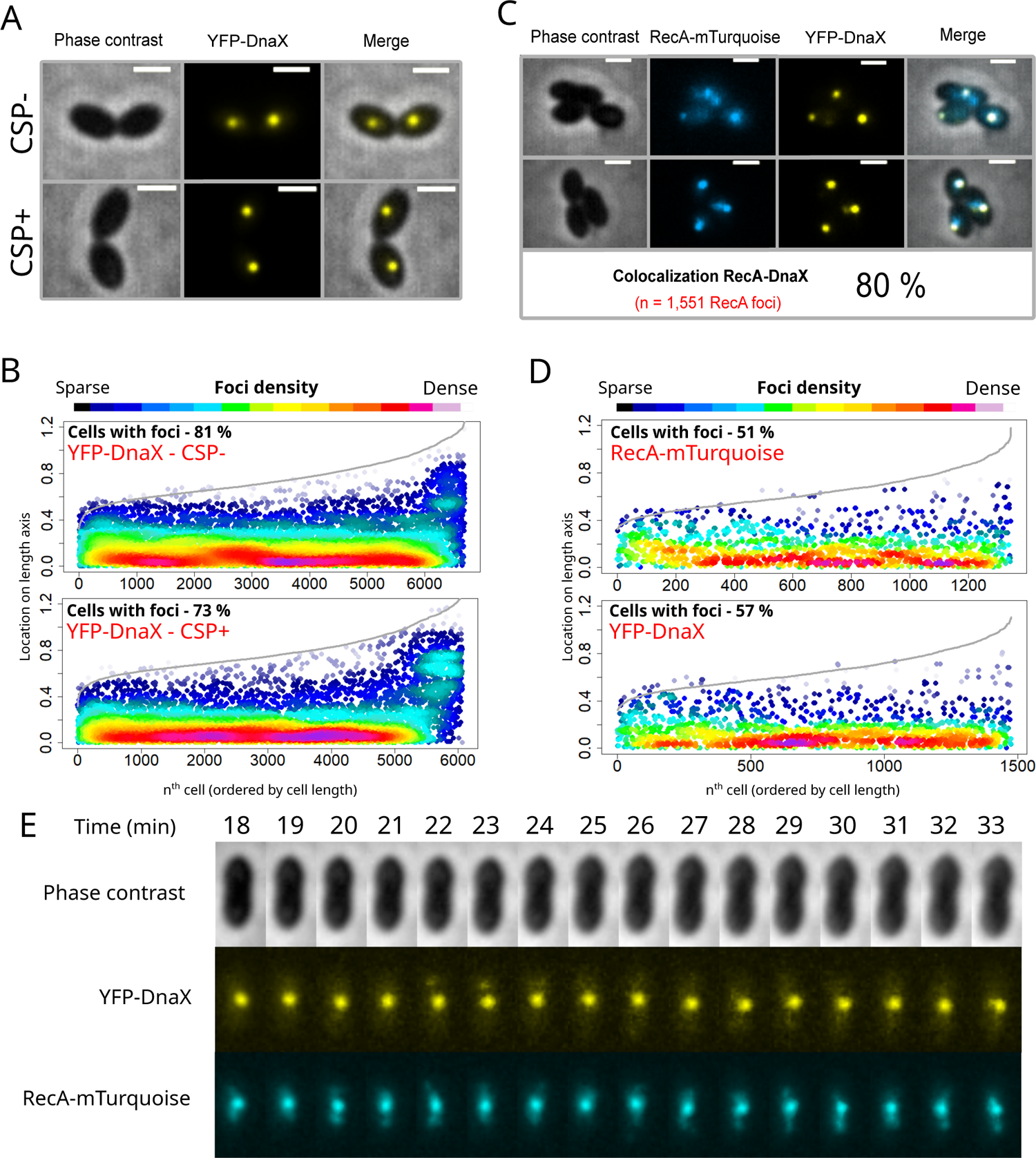
Further exploration of interaction between replication and transformation machineries. (A) Sample microscopy images of cells possessing YFP-DnaX in competence and non-competent cells. Strain used, R4840 (*comC0, ssbB::luc, CEP_M_-yfp-dnaX, CEPII-P_lac_-recA-mturquoise*). (B) Competence mediates a slight reduction in YFP-DnaX foci. Representations as focus density maps as described in Figure 1C. CSP-, 7,911 cells and 6,947 foci analysed. CSP+, 8,425 cells and 6,055 foci analysed. (C) RecA-mTurquoise and YFP-DnaX colocalise in competent cells in the presence of heterologous tDNA (*E. coli* gDNA). Strain used, R4840 (*comC0, ssbB::luc, CEP_M_-yfp-dnaX, CEPII-P_lac_-recA-mturquoise*). 2,443 cells, 1,345 RecA- mTurquoise foci and 1,475 YFP-DnaX foci analysed. (D) Focus density maps of RecA- mTurquoise and YFP-DnaX, as described in Figure 1C. Strain, cell and foci details as in *panel C*. (E) Time-lapse images of RecA-mTurquoise and YFP-DnaX with time representing time after CSP addition (tDNA added at t = 10 min). Strain used as in *panel C*.

**Extended Figure 9:**
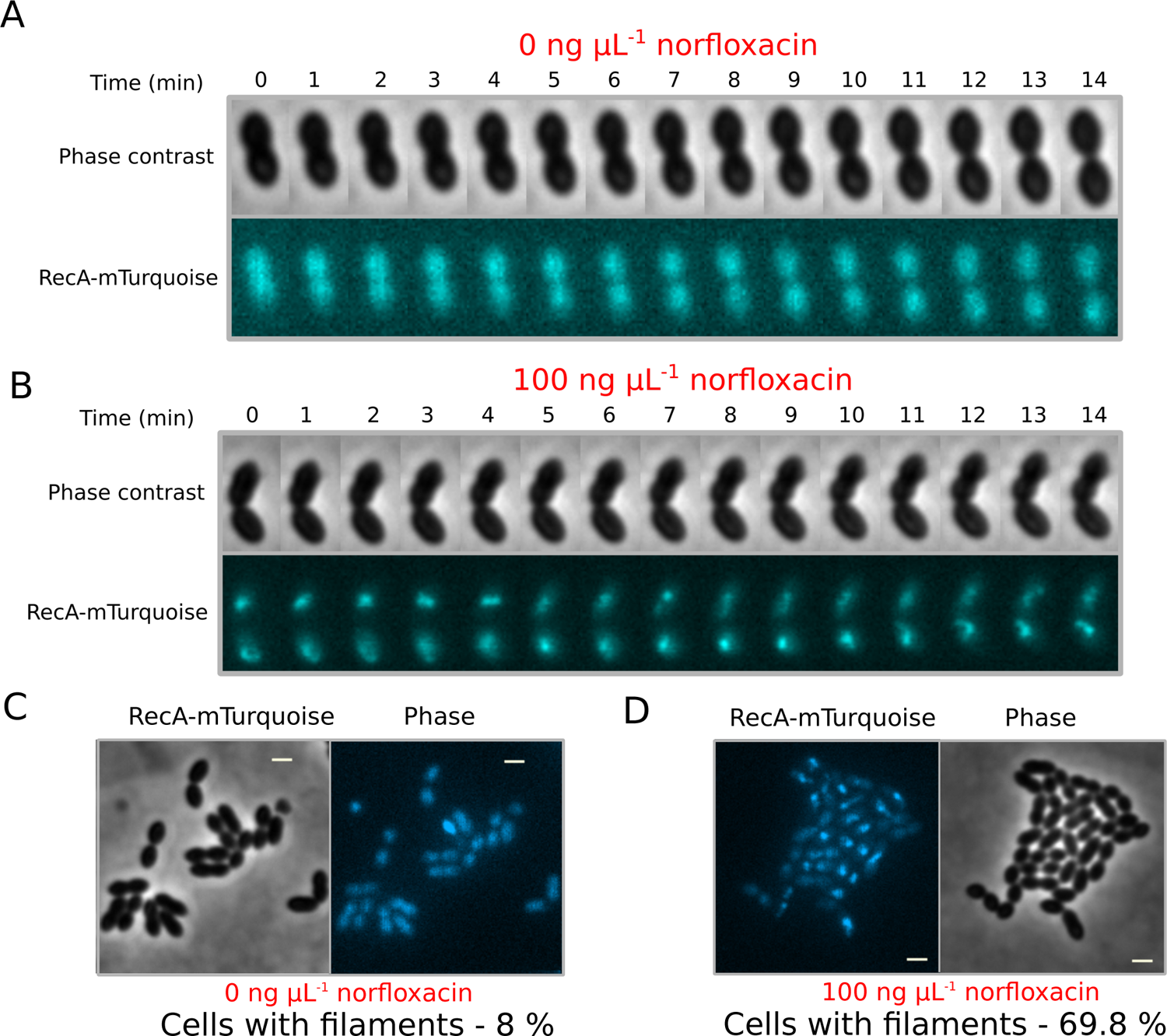
Exploring RecA-mTurquoise filaments and their physiological relevance in genome maintenance. (A) Most RecA/RecA-mTurquoise cells do not display RecA- mTurquoise accumulation in absence of norfloxacin exposure. RecA-mTurquoise observed during time-lapse microscopy of strain R4848 (*comC0, CEPlac-recA-mturquoise*). Images taken every 1 min. (B) Most RecA/RecA-mTurquoise cells display RecA-mTurquoise accumulation into filaments in presence of norfloxacin. RecA-mTurquoise observed during time-lapse microscopy of strain R4848 (*comC0, CEP_lac_-recA-mturquoise*) after 20 min exposure to 100 ng µL^-1^ norfloxacin (MIC 3 ng µL^-1^). Images taken every 1 min. (C) Single image of cells showing lack of RecA-mTurquoise accumulation in absence of norfloxacin exposure. Scale bar, 1 µm. Strain used as in *panel A.* (D) Single image of cells showing RecA-mTurquoise filamentation in almost all cells after norfloxacin exposure. Scale bar, 1 µm. Strain used as in *panel A*.

**Extended Figure 10:**
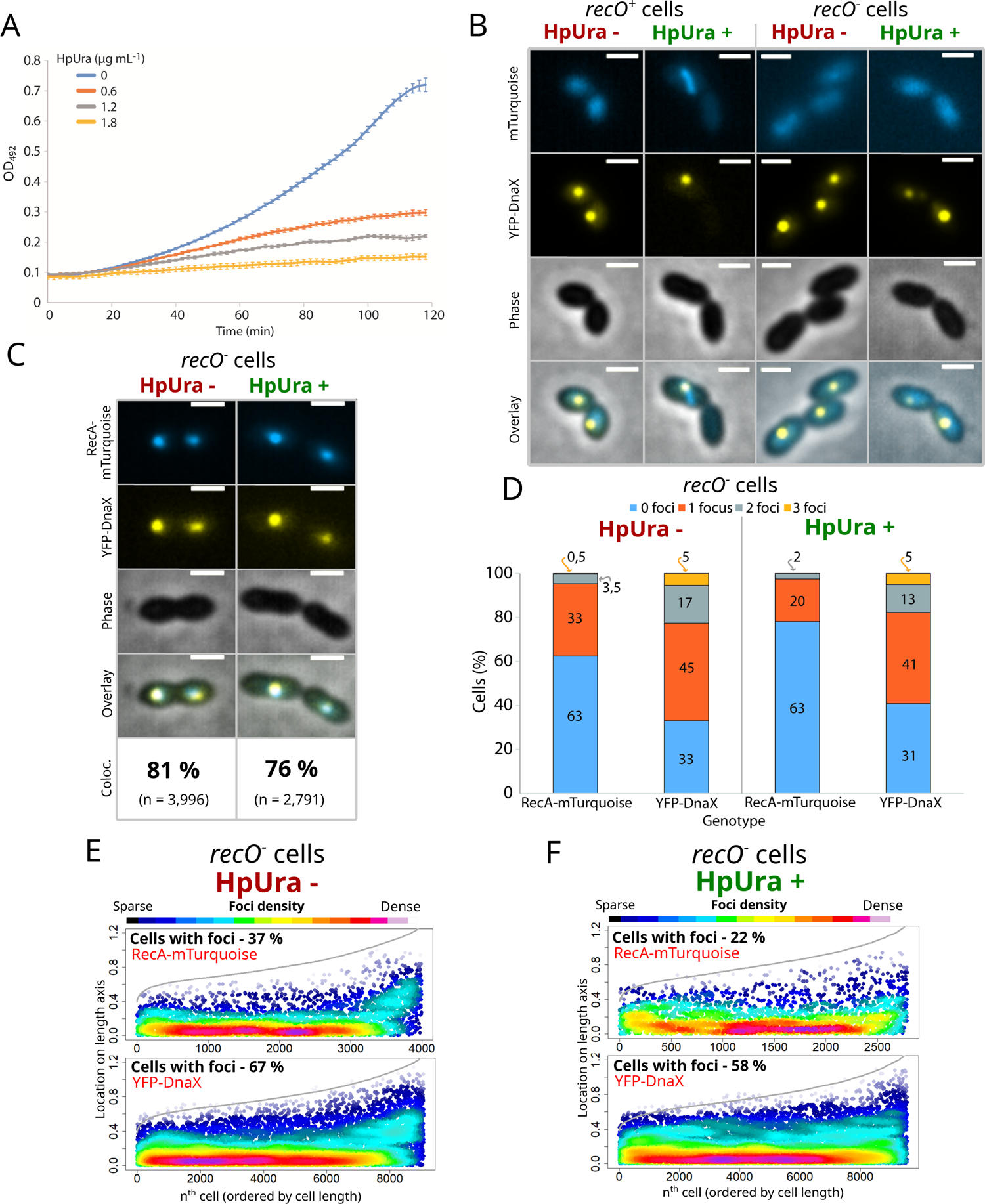
Early HR intermediates of transformation can access stalled replication forks. (A) Growth of pneumococcal cells in presence of varying concentrations of HpUra. Strain used, R1501 (*comC0*). (B) Comparison of RecA/RecA-mTurquoise and YFP-DnaX localisation in non-competent *recO^+/-^* cells exposed to HpUra (1,8 µg mL^-1^) or not. (C) Colocalisation of RecA-mTurquoise and YFP-DnaX foci in competent *recO^-^* cells transformed with homologous tDNA in presence or absence of HpUra. Strain used, R4892 (*comC0, ssbB::luc, CEP_M_-yfp-dnaX, CEPII-P_lac_-recA-mturquoise, recO::spc*). HpUra-, 9,391 cells, 3,996 RecA-mTurquoise foci and 9,072 YFP-DnaX foci analysed. HpUra+, 11,358 cells, 2,791 RecA- mTurquoise foci and 9,555 YFP-DnaX foci analysed. (D) Distribution of RecA-mTurquoise and YFP-DnaX foci per cell. Strain and data used as in *panel C.* (E) Focus density maps of RecA-mTurquoise and YFP-DnaX in transforming *recO^-^* cells in the absence of HpUra. Strain and data used as in *panel C.* (F) Focus density maps of RecA-mTurquoise and YFP-DnaX in transforming *recO^-^* cells in the presence of HpUra. Strain and data used as in *panel C*.

## Movie Legends

Movie 1: Early HR intermediates visualised via DprA-mTurquoise navigate with the dynamic replisome around midcell during transformation of competent pneumococci. Time-lapse microscopy of strain R4631 (*comC0, CEP_M_-yfp-dnaX, CEPII-P_lac_-dprA-mturquoise, dprA::spc*). Images taken at two minute intervals starting 10 min after competence induction and 5 min after DNA addition. Still images from Movie used to make Figure 4D.

Movie 2 – RecA-mTurquoise does not accumulate into filaments in most RecA/RecA- mTurquoise cells in the absence of norfloxacin exposure. Time-lapse microscopy of individual cell of strain R4848 (*comC0, CEP_lac_-recA-mturquoise*) in the absence of norfloxacin. Images taken at 1 min intervals. Still image from movie used in Extended Figure 9A.

Movie 3 – RecA-mTurquoise accumulates into filaments in most RecA/RecA-mTurquoise cells after norfloxacin exposure. Time-lapse microscopy of individual cell of strain R4848 (*comC0, CEP_lac_-recA-mturquoise*) in the presence of norfloxacin (100 ng µL^-1^). Images taken at 1 min intervals. Still image from movie used in Extended Figure 9B.

Movie 4: Early HR intermediates visualised via RecA-mTurquoise navigate with the dynamic replisome around midcell during transformation of competent pneumococci. Time-lapse microscopy of strain R4840 (*comC0, CEP_M_-yfp-dnaX, CEPII-P_lac_-recA-mturquoise*) taken using microfluidics. Images taken at one min intervals starting 5 min after competence induction and immediately upon DNA addition. Still images from Movie used to make Figure 5E.

## Supplementary information

### Supplementary Materials and Methods

#### Protein purification

To purify DprA-GFP, the dprA-gfp sequence was amplified from R3728^38^ using primer pair oALS12 and oALS13; The resulting DNA fragment was digested with *Eco*RI and *Eag*I restriction enzymes and ligated into a pET21 vector digested with the same enzymes to generate the pALS1 plasmid. This plasmid was transformed into *Escherichia coli* Rosetta cells and cells were grown at 37 °C to OD_550_ 0.8 with 0.5 mM IPTG to stimulate DprA-GFP expression. Purification was achieved by sequential passage through three columns as follows; HiTrap Heparin HP 1 mL, gel filtration Superdex 200 Hiload 16/60, HiTrap Q HP 1 mL.

#### In vitro HR assays

HR assays were carried out as follows. 75 nM of DprA or DprA-GFP were incubated at 37 °C for 10 min with varying concentrations of wild type RecA (150, 300, 600 nM) in the presence of 10 nM of Cy3-tagged ovio54 primer (70 nt, fully homologous sequence to pUC18), 10 mM MgOAc and 2 mM ATP. 5 mM of pUC18 plasmid was then added followed by incubation at 37 °C for 10 min. A 1/20 volume of xylene cyanol was added and samples were then denatured by addition of 0.1 % SDS and 10 mM EDTA followed by 3 min incubation at 37°C. Samples were then run on a 1.25 % TBE gel for 60 min at 50 V and DNA was then directly detected on the gel using the Typhoon Trio. Quantification of HR was carried out using MultiGauge software.

#### Plasmid and strain construction

Here we describe how the new plasmids and mutant strains used in this study were generated. Previously published constructs and mutants were simply transferred from published strains by transformation with appropriate selection. The pCJ1 plasmid was generated by removing the MCS from the *pUC57-CEPII_R_-comX* plasmid ^66^. To achieve this, the plasmid was digested with *Eco*RV enzyme, and the insert side recovered. The pUC57 side of the plasmid was amplified with primer pair CJ735-CJ736, each possessing *Eco*RV enzyme sites, removing the MCS site in the process. The insert and PCR were ligated together to generate pCJ1. The pCJ2 plasmid was generated by amplifying a *lacI-P_lac_* PCR fragment from R3833 with primer pair CJ567-CJ730 and a *dprA-mturquoise* PCR fragment from R4062 with primer pair CJ731-CJ595. pCJ1 was digested by *Sal*I and *Kpn*I enyzmes, *lacI-P_lac_* by *Sal*I and *Nco*I enzymes and *dprA-mturquoise* by *Nco*I and *Kpn*I, and the three fragments were ligated together to generate pCJ2. The pCJ3 plasmid was generated by digesting the pMB42 plasmid with the *Xho*I and *Hind*III enzymes to remove *gfp* and ligating in an *mTurquoise* PCR fragment amplified from the R4011 strain^67^ with CJ455-CJ456 primer pair, digested with the same restriction enzymes. The pCJ4 plasmid was generated by amplifying two adjacent DNA fragments by PCR on the R3728 strain^66^ around *dprA-gfp* construct using primer pairs CJ391-CJ465 and CJ466- CJ378 respectively. The three base mutations required to alter *gfp* to *yfp* present in both primers CJ465 and CJ466. Splicing overlap extension (SOE) PCR on these two fragments with the CJ391-CJ378 primer pair generated a DNA fragment with the *yfp* mutation. This DNA fragment was transformed without selection into R3728 ^66^ with a 3 h 30 min phenotypic expression phase in liquid culture to introduce the *yfp* mutation, and positive clones were determined by PCR amplification with the CJ391-CJ378 primer pair and sequencing with the CJ378 primer. The pCJ5 plasmid was generated by digesting the pMB42 plasmid ^66^ with the *Eco*RI and *Xho*I enzymes to remove *‘dprA* and ligating in a *‘recA* PCR fragment amplified from the R1501 strain with CJ764-CJ765 primer pair, digested with the same restriction enzymes. The pCJ6 plasmid was generated by amplifying a PCR fragment consisting of the P*_lac_* promoter and upstream *lacI* gene from R4261 ^66^ using primer pair CJ567-CJ615 and digesting it with *Sal*I and *Nco*I enzymes. A *dprA^QNQ^-gfp* DNA fragment was amplified from R4046 ^66^ using primer pair CJ411-CJ616 and digested with *Nco*I and *Bam*HI enzymes. The *pCEP_R_-luc* plasmid was digested with *Sal*I and *Bam*HI enzymes and these three fragments were ligated together to generate pCJ6. The pCJ7 plasmid was generated in the same manner but with an amplification of a *dprA^AR^-gfp* DNA fragment was amplified from R4047 ^66^ using primer pair CJ411-CJ616. The R2546 strain (*comC0, CEP_X_-gfp*) was constructed by transforming R1501 with the pCN35 plasmid ^68^ and selecting for kanamycin resistance. The R3406 strain (*comC0, ssbB-luc, CEP_M_- yfp-dnaX*) was generated by making four DNA fragments by PCR; a fragment of the upstream CEP platform sequence from pCEP ^69^ with primer pair OVK53-OVK54; the *yfp* sequence from R4404 with primer pair OVK55-OVK56; the *dnaX* sequence from R1501 with primer pair OVK61-OVK62 and the downstream CEP platform sequence from pCEP ^69^ with primer pair OVK57-OVK73. A SOE PCR fragment was generated using these four fragments with primer pair OVK53-OVK73, and this was transformed into R1502, with transformants selected with kanamycin. The R4062 strain (*comC0, dprA-mturquoise*) was generated by transforming R1501 with the pCJ3 plasmid and selecting for spectinomycin resistance. The R4400 strain (*comC0, CEP_lac_-dprA-gfp, ssbB::cat*) was generated by transforming R4262 with genomic DNA from the R4812 strain and selecting for kanamycin resistance. The R4401 strain (*comC0, CEP_lac_-dprA-gfp, comEC::ery*) was generated by transforming R4262 with genomic DNA from the R2586 strain ^37^ and selecting for erythromycin resistance. The R4404 strain (*comC0, dprA- yfp*) was generated by transforming R1501 with the pCJ4 plasmid and selecting for spectinomycin resistance. The R4412 strain (*comC0, CEP_lac_-dprA^QNQ^-gfp*) was generated by transforming R1501 with pCJ6 and selecting transformants with kanamycin. The R4413 strain (*comC0, CEP_lac_-dprA^AR^-gfp*) was generated by transforming R1501 with pCJ7 and selecting transformants with kanamycin. The R4415 strain (*comC0, CEP_lac_-dprA^QNQ^-gfp, dprA::spc*) was generated by transforming R4412 with genomic DNA from the R751 strain ^70^ and selecting for spectinomycin resistance. The R4416 strain (*comC0, CEP_lac_-dprA^AR^-gfp, dprA::spc*) was generated by transforming R4413 with genomic DNA from the R751 strain ^70^ and selecting for spectinomycin resistance. The R4429 strain (*comC0, CEP_lac_-dprA-gfp, dprA::spc, recA::cat*) was generated by transforming R4262 with genomic DNA from the R209 strain ^71^ in presence of 50 µM IPTG and selecting for chloramphenicol resistance. To generate the R4618 strain (*comC0, CEP_lac_-dprA-gfp, dprA::spc, comEC::trim, recA::cat*), a *comEC*::*trim* DNA fragment was created by initial amplification of the regions upstream and downstream of the *comEC* gene using primer pairs CJ720-721 and CJ724-725 and R1501 gDNA as template. The trimethoprim resistance cassette was amplified using the primer pair CJ722-723 and the R4107 strain ^66^ as template. SOE PCR on these three fragments with the primer pair CJ720-725 generated a DNA fragment with the *comEC* gene replaced with the trimethoprim resistance cassette, which was co-transformed into R4262 with a *recA::cat* DNA fragment amplified from R209 ^71^ using primer pair CJ726-CJ727. Transformants were selected with trimethoprim and chloramphenicol to integrate both *comEC::trim* and *recA::cat* at the same time, since both abrogate transformation. To generate the R4625 strain (*comC0, CEP_lac_-dprA-gfp, dprA::spc, radA::trim*), a *radA*::*trim* DNA fragment was created by initial amplification of the regions upstream and downstream of the *radA* gene using primer pairs CJ748-CJ749 and CJ752-oIM58 and R1501 gDNA as template. The trimethoprim resistance cassette was amplified using the primer pair CJ750-751 and the R4107 strain ^66^ as template. SOE PCR on these three fragments with the primer pair CJ748-oIM58 generated a DNA fragment with the *radA* gene replaced with the trimethoprim resistance cassette, which was transformed into R4262 ^66^ in presence of 50 µM IPTG and transformants were selected with trimethoprim. To generate the R4626 strain (*comC0, ssbB-luc, CEP_M_-yfp-dnaX, CEPII_lac_-dprA-mTurquoise*), R3406 was transformed with the pCJ2 plasmid, and transformants were selected with erythromycin. To generate the R4631 strain (*comC0, ssbB-luc, CEP_M_-yfp-dnaX, CEPII_lac_-dprA-mTurquoise, dprA::spc*), R4626 was transformed with genomic DNA from strain R751 and transformants were selected with spectinomycin. To generate strain R4664, a fragment of *CEP_lac_* was amplified using primer pair CJ588-CJ680 and R3833 as template, and the *recA* gene was amplified using primer pair CJ681- CJ682 and R1501 as template. The pCEPlac-dprA-gfp plasmid was digested with *Sal*I and *Bam*HI restriction enzymes, while the DNA fragments were digested with *Sal*I/*Nco*I and *Nco*I/*Bam*HI respectively. These three fragments were ligated together and transformed into R1501, with transformants selected with kanamycin. To generate strain R4712 (*comC0, recA- mTurquoise*), R1501 was transformed with pCJ5 and transformants were selected with spectinomycin. To generate strain R4716 (*comC0, CEP_lac_-dprA-gfp, recA-mTurquoise*), R4261^66^ was transformed with pCJ5 and transformants were selected with spectinomycin. To generate strain R4731 (*comC0, CEP_lac_-dprA-yfp, recA-mTurquoise*), two adjacent DNA fragments were amplified by PCR on the R4262 strain ^66^ around *CEP_lac_*-*dprA-gfp* construct using primer pairs CJ114-CJ465 and CJ466-kan1 respectively. The three base mutations required to alter *gfp* to *yfp* present in both primers CJ465 and CJ466. SOE PCR on these two fragments with the CJ114-kan1 primer pair generated a DNA fragment with the *yfp* mutation. This DNA fragment was transformed without selection into R4716 with a 3 h 30 min phenotypic expression phase in liquid culture to introduce the *yfp* mutation, and positive clones were determined by PCR amplification with the CJ114-kan1 primer pair and sequencing with the CJ114 primer. To generate the R4742 strain (*comC0, CEP_lac_-dprA-yfp, recA-mTurquoise, dprA::trim)*, a *dprA*::*trim* DNA fragment was created by initial amplification of the regions upstream and downstream of the *dprA* gene using primer pairs CJ373-CJ770 and CJ773-CJ378 and R1501 gDNA as template. The trimethoprim resistance cassette was amplified using the primer pair CJ771-CJ772 and the R4107 strain ^66^ as template. SOE PCR on these three fragments with the primer pair CJ373-CJ378 generated a DNA fragment with the *dprA* gene replaced with the trimethoprim resistance cassette, which was transformed into R4731 and transformants were selected with trimethoprim. The R4812 strain (*comC0, ssbB::cat*) was generated by transforming R2294 ^43^ with the *pEMcat* plasmid and transformants were selected with chloramphenicol. To generate strain R4840, the regions upstream and downstream of *dprA-mturquoise* in the CEPII platform were amplified from R4631 using primer pairs CJ662-CJ793 and CJ667-CJ796 respectively, and *recA-mturquoise* was amplified from R4712 using primer pair CJ794-CJ795. SOE PCR using there three fragments and primer pair CJ662-CJ667 generated a *CEPII-P_lac_-recA-mturquoise* DNA fragment which was transformed into R3406, with transformants selected with erythromycin. To generate strain R4848 (*comC0, CEP_lac_-recA-mturquoise*), 5’ and 3’ fragments of *CEP_lac_* were amplified from R4262 with primer pairs CJ574-CJ799 and CJ802-CJ575 respectively, and *recA- mturquoise* was amplified from R4712 using primer pair CJ800-CJ801. SOE PCR with these DNA fragments and primer pair CJ574-CJ575 generated a *CEP_lac_-recA-mturquoise* fragment which was transformed into R1501 and transformants were selected with kanamycin. To generate strain R4849 (*comC0, dprA-lgbit*), 5’ and 3’ fragments of *dprA* were amplified from R1501 with primer pairs CJ689-CJ690 and CJ693-CJ694, and the *lgbit* tag with appropriate linker and over hang sequences for SOE PCR was synthesized based on previously-published sequence optimized for the pneumococcus using gBlocks (Intergrated DNA technologies). SOE PCR with these DNA fragments and primer pair CJ689-CJ694 generated a *dprA-lgbit* fragment which was transformed into R1501 without selection and transformants were screened by PCR for integration using primer pair CJ689-CJ694. To generate strains R4851 (*comC0, CEP_lac_- recA-mturquoise, dprA::spc*), R4848 was transformed with chromosomal DNA from R751 (*rpsL41, dprA::spc*) ^70^ and transformants were selected with spectinomycin. To generate strain R4856, a DNA fragment containing the *smbit* tag fused to the 3’ end of the *dnaX* gene with a linker, flanked by 5’ and 3’ sequences of *dnaX* was generated using gBlocks (Integrated DNA technologies). 5’ and 3’ fragments of *dnaX* were amplified from R1501 with primer pairs CJ809-CJ810 and CJ813-CJ814, and SOE PCR using these three DNA fragments and primer pair CJ809-CJ814 generated a *dnaX-smbit* DNA fragment which was transformed into R1501 without selection and transformants were screened by PCR for integration using primer pair CJ811-CJ812. To generate strain R4857 (*comC0, ΔrecA::trim*), upstream and downstream sequences around the *recA* gene were amplified using primer pairs CJ829-CJ830 and CJ833- CJ808 respectively, and the TrimR resistance cassette was amplified from strain R4107 ^66^ using primer pair CJ831-CJ832. A *ΔrecA::trim* DNA fragment was generated by SOE PCR using these three DNA fragments and transformed into R1501, with transformants selected with trimethoprim. To generate strain R4858 (*comC0, dprA-lgbit, CEP_lac_-dprA-smbit*), two DNA fragments containing upstream and downstream sequences around the 3’ end of *dprA* in *CEP_lac_-dprA* were amplified from strain R4262 using primer pairs CJ574-CJ827 and CJ575- CJ828 respectively, where primers CJ827 and CJ828 include the *linker-smbit* sequence. SOE PCR using these two DNA fragments generated a *CEP_lac_-dprA-smbit* fragment, which was transformed into R4849 and transformants were selected with kanamycin. To generate strain R4859 (*comC0, dprA-lgbit, dnaX-smbit, hexA::ermAM*), R4856 cells were transformed with genomic DNA from strain R246 (*hexA::ermAM*) and transformants were selected with erythromycin. To generate strain R4861, R4859 cells were transformed with a *dprA^AR^* PCR fragment amplified from strain R2585 ^24^ using primer pair CJ311-CJ391, and transformants were screened by PCR and sequencing (Eurofins MWG) for insertion of the two independent mutations conferring the dprA^AR^ phenotype ^24^.

#### Time-lapse microfluidics experiments

Time-lapse microfluidics experiments were carried out using a CellASIC ONIX Microfluidic platform and B04A microfluidic plates (Merck-Millipore, Billerica, MA, U.S.A) as previously described ^72^, with modifications. Briefly, exponentially growing cultures (OD_550_ 0,3) of R4840 (*comC0, CEP_M_-yfp-dnaX, CEPII-P_lac_-recA-mturquoise*) were diluted 50-fold in C+Y medium (supplemented with 300 U/mL catalase, 0.3 % maltose and 50 µM IPTG) and incubated at 37 °C to an OD_550_ of 0,1. Cells were then loaded into the microfluidic chamber and maintained at 37 °C in a thermostated chamber with a constant flow rate of 0,3 µL/h (0,25 psi). Competence induction was achieved by injecting CSP (1 µg mL^-1^ in C+Y medium with catalase, maltose and IPTG) for 3 min at 6 psi. DNA (250 ng µL ^-1^, diluted in C+Y medium with catalase, maltose and IPTG) was then injected for 6 min at 3 psi, followed by 1 h at 0.25 psi. Images were captured every minute throughout using the same microscope set-up as described above, with a thermostated chamber at 37 °C.

#### Sensitivity to DNA damage assays

Survival assays were performed as previously described ^56^, with modifications. Briefly, cells were grown to OD_550_ 0.1 in C+Y medium (with 50 µM IPTG where appropriate) before serial dilution and spotting of 10 µL volumes onto pre-dried plates containing 0.02 % MMS and 50 µM IPTG where appropriate. After spot drying, plates were incubated overnight at 37°C in a bell jar with Anaerocult A (Merck) to promote anaerobic conditions.

#### Fluorescent DNA microscopy experiments

Two independent DNA fragments (*rpsL1* and *radA::spc*) were used to test internalization of transforming DNA, *rpsL1*, conferring streptomycin resistant via point mutation, or *radA::spc*, conferring spectinomycin resistance by integration of a heterologous cassette. A 2,008 bp DNA fragment containing *rpsL1* was amplified from R2980 (*dpnMAB, rpsL1*) using primer pair MB83-MB84. A 4,649 bp DNA fragment containing *radA::spc* was amplified from R3255 (*dpnMAB, hexA::ermAM, radA::spc, ssbB::kan*) using primer pair CJ338- CJ368. Strains possessing the *Dpn*II restriction system were used as templates to allow template removal by *Dpn*I digestion. PCR fragments were labelled with either fluorescein, Dylight 550 or Dylight 650. For fluorescein labelling, 1 µL of 1 mM fluorescein-12-dUTP (Thermo Fisher Scientific), 2 µL dNTP mix (1 mM dATP, dCTP, and dGTP and 0.5 mM dTTP (Thermo Fisher Scientific)), 0.5 µL DreamTaq DNA polymerase (Thermo Fisher Scientific), 5 µL DreamTaq buffer, 1 µM of each primer and 2 µL of genomic DNA were used in a total reaction mixture volume of 50 µL. The reaction conditions for Dylight labelling were the same as for fluorescein, but with 1 µL of dNTP mixture (10 mM dGTP, dCTP, and dATP and 5 mM dTTP and aminoallyl-dUTP (Thermo Fisher Scientific)). After labelling, samples were protected from light throughout. Samples were then mixed 4:1 with Dylight 550 or 650 (10 mg mL^-1^, Thermo Fisher Scientific) and incubated at room temperature for 3 hours. Samples were then incubated for 2 h with 0.5 µL *Dpn*I (20 U µL^-1^, FastDigest, Thermo Fisher Scientific) per 50 µL PCR sample. Label incorporation was calculated using a NanoDrop (Thermo Fisher Scientific) as follows: fluorescein, 0,3-1,6 pmol µL^-1^; Dylight 550 and 650, 0,9-4,4 pmol µL^-1^. Microscopy images were captured as described above using FITC (fluorescein), Cy3 (Dylight 550) and Cy5 (Dylight 650) filters respectively.

### Supplementary Results

#### Exploring the visualization of fluorescent tDNA during pneumococcal transformation

In this study, fluorescent fusions of DprA and RecA were used to visualize the early HR intermediates in actively growing pneumococci. The other main actor of early HR intermediates is ssDNA, and the possibility of visualizing fluorescent transforming ssDNA in *B. subtilis* and *S. pneumoniae* has previously been explored, showing fluorescent foci on cells after DNase I treatment ^47^. However, a further study in *B. subtilis* showed that resistance to DNase I does not necessarily indicate entry into the cytoplasm, but rather the periplasm ^19^. In light of this, the potential of using such fluorescent DNA to visualize early HR intermediates during transformation was explored in *S. pneumoniae*. To begin, the transformation efficiency of labelled DNA fragments possessing an *rpsL41* point mutation ^73^ was compared to unlabelled controls with dUTP/dTTP mix or dTTP alone. Results showed that the dUTP/dTTP unlabelled mix showed reduced transformation efficiency compared to dTTP alone, while labelled DNA fragments showed similar or slightly reduced transformation efficiency compared to the dUTP/dTTP unlabelled mix (Extended Figure 7A). This suggested that fluorescent DNA could be internalised, however, internalization and integration of short unlabelled fragments around the point mutation could not be excluded. Fluorescence microscopy on competent cells transforming with labelled DNA fragments showed distinct foci associated to a minority of cells (Extended Figure 7B). To further explore whether fluorescently labelled DNA could be internalized by competent cells, transformation experiments were carried out using DNA fragments possessing a point mutation as above (*rpsL1*, otherwise homologous DNA) or a heterologous antibiotic resistance cassette flanked by homologous sequences (*radA::spc*), labelled or not with Dylight 500 or 650, at varying DNA concentrations. Unlike a point mutation, integration of a heterologous cassette requires transfer of the entire cassette plus flanking sequences, making the presence of entirely unlabelled fragments of transformable DNA much less likely. Results show that while reducing the concentration of *rpsL1* DNA below saturating levels did not alter the ratio between transformation efficiency of labelled and unlabelled DNA donors, reducing the concentration of *radA::spc* donor DNA specifically reduced the transformation efficiency of labelled DNA, increasing the ratio of transformation efficiency between labelled and unlabelled DNA (Extended Figure 7CD). This result suggests that high levels of fluorescent labelling negatively impacted the transformation of a donor DNA molecule, but that nonetheless, less labelled donor DNA can be internalized and integrated into the recipient chromosome by transformation. To explore whether it was possible to visualize this subpopulation of transforming DNA, fluorescence microscopy was carried out on wildtype cells in the presence or absence of CSP, as well as in several transformasome mutant strains. Results show that in wildtype cells, although foci are observed in non-competent cells, competence specific foci are observed (Extended Figure 7E). The absence of the pilus (*comGA^-^*), the EndA nuclease (*endA^-^*) or the transformation-dedicated recombinase loader (*dprA^-^*) did not alter the number of competent cells possessing foci despite being key for capture, processing and protection of transforming DNA, respectively. However, a reducing in cells possessing foci was observed in the absence of the DNA transformation pore (*comEC^-^*), while removing the DNA receptor (*comEA^-^*) reduced the number of cells possessing foci to non-competent levels (Extended Figure 7E). The localization of foci did not vary significantly in all of these strains (Extended Figure 7F). In conclusion, although it appears DNA molecules possessing fewer fluorescent tags can be transformed, visualisation of these is not possible due to the pollution from DNA molecules with greater numbers of fluorescent tags, which are not transformable and thus remain on the outside of the cell.

